# Viscoelastic high-molecular-weight hyaluronic acid hydrogels support rapid glioblastoma cell invasion with leader-follower dynamics

**DOI:** 10.1101/2024.04.04.588167

**Authors:** Emily M. Carvalho, Erika A. Ding, Atul Saha, Anna Weldy, Peter-James H. Zushin, Andreas Stahl, Manish K. Aghi, Sanjay Kumar

**Affiliations:** Department of Chemical and Biomolecular Engineering, University of California, Berkeley, CA 94720, USA; Department of Neurosurgery, University of California, San Francisco, CA 94158, USA; Department of Nutritional Sciences and Toxicology, University of California, Berkeley 94720, USA; Department of Bioengineering, University of California, Berkeley, CA 94720, USA; Department of Bioengineering and Therapeutic Sciences, University of California, San Francisco, CA 94158, USA

**Keywords:** hyaluronic acid (HA), high-molecular-weight (HMW), viscoelastic, stress-relaxing (SR), glioblastoma (GBM)

## Abstract

Hyaluronic acid (HA), the primary component of brain extracellular matrix, is increasingly used to model neuropathological processes, including glioblastoma (GBM) tumor invasion. While elastic hydrogels based on crosslinked low-molecular-weight (LMW) HA are widely exploited for this purpose and have proven valuable for discovery and screening, brain tissue is both viscoelastic and rich in high-MW (HMW) HA, and it remains unclear how these differences influence invasion. To address this question, hydrogels comprised of either HMW (1.5 MDa) or LMW (60 kDa) HA are introduced, characterized, and applied in GBM invasion studies. Unlike LMW HA hydrogels, HMW HA hydrogels relax stresses quickly, to a similar extent as brain tissue, and to a greater extent than many conventional HA-based scaffolds. GBM cells implanted within HMW HA hydrogels invade much more rapidly than in their LMW HA counterparts and exhibit distinct leader-follower dynamics. Leader cells adopt dendritic morphologies, similar to invasive GBM cells observed in vivo. Transcriptomic, pharmacologic, and imaging studies suggest that leader cells exploit hyaluronidase, an enzyme strongly enriched in human GBMs, to prime a path for followers. This study offers new insight into how HA viscoelastic properties drive invasion and argues for the use of highly stress-relaxing materials to model GBM.

## 1. Introduction

Mechanical properties of the extracellular matrix (ECM) are widely recognized to influence cell behavior, including morphology, motility, proliferation, and differentiation.^1–4^ While attention in the field initially focused on the effects of ECM elastic (storage) properties, most solid tissues are viscoelastic,^5^ meaning that they can both store and dissipate applied stresses. Growing recognition of the biological importance of viscoelasticity has motivated a significant effort to develop biomaterial culture scaffolds with tunable viscoelastic properties. Collagen and alginate hydrogels are perhaps the most widely used systems to study the effects of viscoelasticity, where the noncovalent crosslinking within these hydrogels allows stress relaxation through the breaking, reformation, and rearrangement of crosslinks.^6^ A variety of strategies have been explored to tune viscoelastic properties within these systems. In one example, static and dynamic covalent bonds were added to an interpenetrating network (IPN) of collagen and hyaluronic acid (HA) to reduce its stress relaxation rate,^7^ and in another, PEG spacers were added to alginate to increase its stress relaxation rate.^8^ Systematic variation of viscoelastic properties within these networks can affect cell behavior in powerful and surprising ways. For mesenchymal stem cell (MSC) fate commitment, increasing viscous properties in a stiff matrix promotes greater MSC osteogenic differentiation whereas reducing viscous properties in a soft matrix promotes adipogenic differentiation.^5^

Careful consideration of ECM viscous properties is particularly important in brain tissue, which is ∼80% water^9^ and exquisitely sensitive to external shear forces,^10^ internal dynamics of blood and cerebrospinal fluid flow,^11^ and changes in intracranial pressure.^12^ For example, the viscoelasticity of brain tissue is increasingly understood to play important roles in the propagation of stresses in traumatic brain injury (TBI),^13^ which has in turn spurred efforts to incorporate viscoelastic properties into therapeutic materials for healing brain lesions associated with TBI and other pathologies.^14^ Two distinguishing features that contribute to the highly viscous nature of brain ECM are its relative lack of fibrous structures (e.g. collagen) and its enrichment in hyaluronic acid (HA), a non-sulfated glycosanimoglycan.^15^ In addition to its structural and mechanical roles, HA serves a series of biological functions, from engaging with cell surface receptors such as CD44, to undergoing remodeling processes via cell surface enzymes capable of HA synthesis (hyaluronan synthases (HAS)) and degradation (hyaluronidases (HYAL)) (**Figure 1a**).^15^ HA plays an especially important role in the progression of the deadly and diffuse brain cancer glioblastoma (GBM). GBM is strongly defined by invasion into the HA-rich brain parenchyma, where tumor cells escape surgical resection, become inaccessible to chemotherapy, and can seed secondary tumors.^16,17^ It has been shown that HA binding and remodeling proteins contribute to and are biomarkers for disease severity.^18–21^

**Figure 1.**
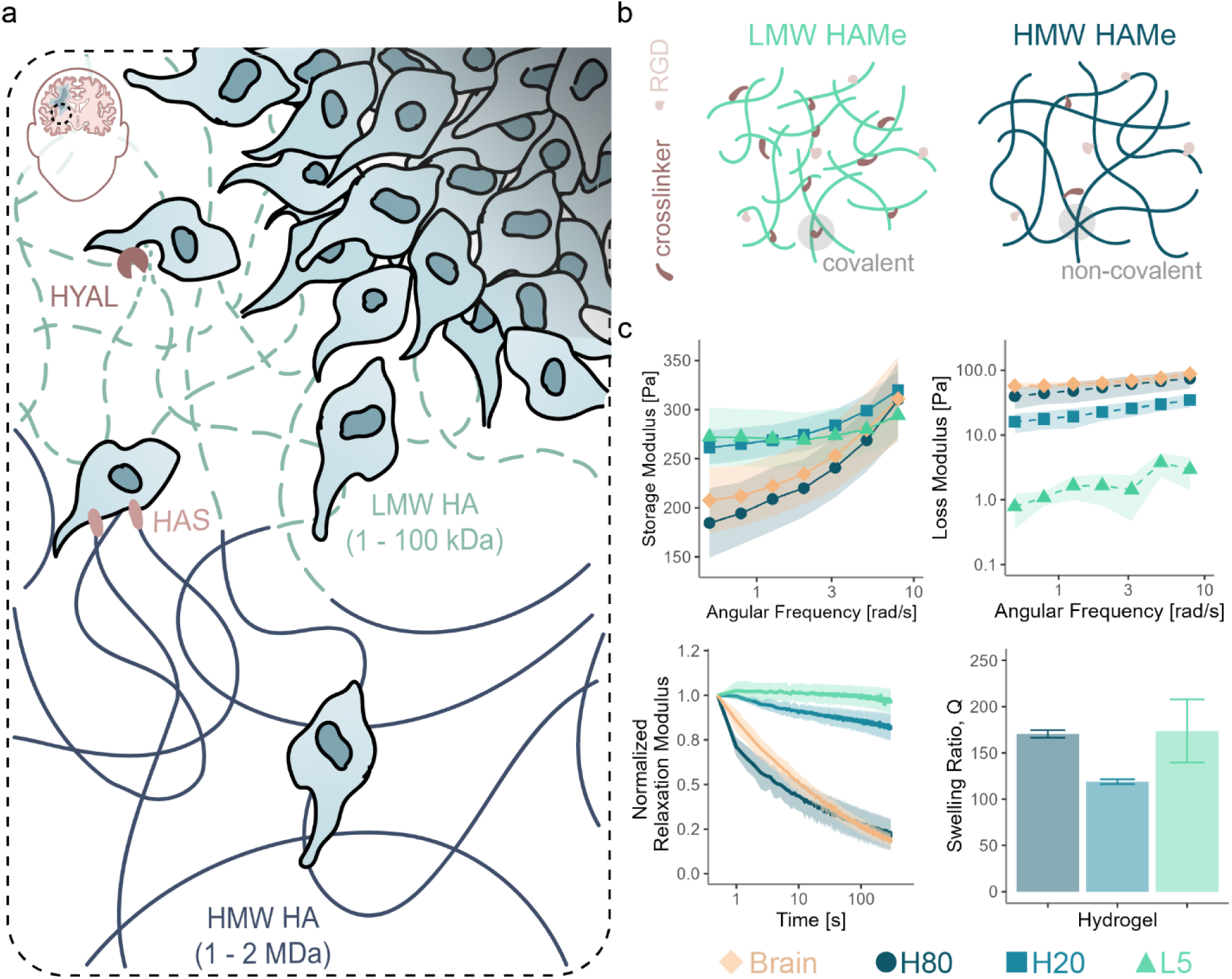
Stress relaxing materials. **a.** Schematic of the GBM microenvironment in HA-rich brain **b.** Schematic of HMW and LMW HA hydrogels **c.** Material tests of HA hydrogels and healthy mouse brain tissue: rheological oscillation tests at 0.5% strain (top; n=3-5), rheological stress relaxation tests with 15% strain over 300s (bottom left; n=3-5), and swelling tests (bottom right; n=3).

The importance of HA in GBM progression has spurred efforts to develop HA-based matrix platforms to model GBM invasion in vitro.^4,22,23^ However, it is not well understood how HA viscoelastic properties contribute to GBM invasion. While HA in the brain is comprised of high-molecular-weight (HMW) chains crosslinked noncovalently with large tenascin proteins decorated with lectins,^15^ HA hydrogels are typically fabricated *in vitro* by chemically modifying the backbone of low-molecular-weight (LMW) HA chains to introduce functional groups that support fast and efficient conjugation of crosslinkers. The crosslinking chemistries traditionally used in HA hydrogels (e.g. Michael addition alkyne-azide cycloaddition) are based on static, covalent bonds that store but do not dissipate applied stresses. Recent efforts have sought to introduce viscoelastic properties into HA hydrogels^24–27^ or HA-based interpenetrating networks (IPNs)^7,28^ through the use of dynamic crosslinks that transiently break to relax stress and then reform. While these systems have begun to lend important insight into how ECM viscous properties modulate cell behavior, it has proven challenging to match the stress relaxation characteristics of hydrogel models to brain tissue. For example, HA hydrogels based on Cucurbit[7]uril host crosslinks (K_eq_ = 10^-6^ – 10^-2^)^29^ relax stresses faster than brain tissue, whereas HA hydrogels based on hydrazone crosslinks (K_eq_ = 10^3^-10^5^)^30^ do so more slowly than brain tissue.

As noted above, HA hydrogel formulations typically employ LMW HA species (∼60 kDa), in part because the low viscosity of LMW HA solutions facilitates synthesis, assembly, and use in cell culture. However, HMW (∼1.5 MDa) HA formulations could create access to a wider range of viscoelastic properties, in part because the longer chains could themselves contribute to stress relaxation through greater chain entanglement, deformation, and relaxation. In this study we explore the use of HMW HA-based hydrogels as viscoelastic culture platforms for modeling GBM invasion. We introduce and characterize a set of HA-based hydrogels that exhibit different extents of stress relaxation, with the most stress-relaxing HMW hydrogel mimicking the stress relaxation properties of brain tissue. We find that increasing HA stress relaxation leads to faster GBM cell invasion, which occurs through a distinct leader-follower mechanism resembling GBM invasion in vivo. Next-generation sequencing, pharmacologic studies, and microscopy suggest that this dynamic depends on the ability of the leader cells to enzymatically remodel the surrounding matrix via hyaluronidases, facilitating the advance of follower cells.

## 2. Results

### 2.1 Polymer entanglements facilitate HMW HA hydrogel stress relaxation

Our first goal was to design HMW HA hydrogels that relax stresses to the same extent and speed as brain tissue. To first address the technical barrier of using a highly viscous HMW polymer, we introduced a number of modifications, including the use of more dilute HA solutions, centrifugation to remove bubbles, and application of positive displacement pipettes to draw accurate volumes and mix solutions (see Methods). We reasoned that HMW HA solutions may be inherently viscoelastic due to entanglement of the constituent polymer chains, which are presumed both to store stresses by straining entanglement-based crosslinks and dissipate stresses through chain sliding and rearrangement. These effects are expected to be both frequency-and concentration-dependent.^31^ We therefore created a series of HMW (1.5 MDa) HA solutions of varying monomer concentration (10 – 40 mg/mL) (**Figure S1a-b**) and measured their rheological properties. As expected, we observed solid-like properties at high frequencies, where the storage modulus is higher than the loss modulus (**Figure S1b**). For 40 mg/mL HA solutions, the storage modulus exceeded the loss modulus at all measured frequencies, while for more dilute HA concentrations the loss modulus exceeded the storage modulus at low frequency, with the “crossover frequency” increasing with decreasing HA concentration (**Figure S1b**). These observations are consistent with a picture in which more concentrated HMW HA solutions create more entanglements, which store stress at high frequencies, and thus behave more like a solid. At lower frequencies these noncovalent crosslinks dissociate, giving rise to more liquid-like behavior. Direct measurement of stress-relaxation properties in the time domain revealed that when a step strain of 15% is applied, all solutions fully relaxed to 0% stress (or in other terms, have a normalized relaxation modulus of 0) within 5 minutes, again consistent with a mechanism in which chain entanglements untangle and rearrange to dissipate the imposed stress (**Figure S1b**). Throughout the paper we will refer to the extent to which stress is reduced (or 1-normalized relaxation modulus) at this time point as % stress relaxing (% SR). Consistent with the frequency sweep measurements, relaxation time increased with HA concentration (**Figure S1b**), presumably due to the time required for additional network rearrangements.

Although our HMW HA solutions are formally solid-like, particularly at the highest HA concentration, they are expected to be poorly suited to long-term cell culture, where swelling and chain diffusion would disintegrate the network. Therefore, we next introduced modest amounts of more permanent crosslinks. In brain tissue these crosslinks are provided by HA-binding proteins (e.g. tenascins), which multivalently and specifically ligate HA chains to stabilize the network.^15^ To simulate these linkages in vitro, we employed Michael Addition-based thioester crosslinks as we and others have done previously to assemble LMW HA hydrogels.^4^ Building from our previously described protocols, we methacrylated our HMW HA polymers (**Figure S1c**) and then crosslinked them into a hydrogel with dithiol peptides, whose sequences can be customized to facilitate protease digestion. While our HMW HA networks became increasingly elastic in nature with the addition of many covalent crosslinks, we could identify a range of covalent crosslink densities (0.0 – 0.05 thiol: monomer ratio (1.676 mM of crosslinker) for a 30 mg/mL HA network) that supported a range of stress relaxation capabilities (∼100% to ∼0% stress relaxation) (**Figure S1d**). Increasing covalent crosslink density also increases the storage modulus over a range of frequencies (0.5 – 50 rad/s), with the most dramatic increases observed at the lowest frequencies (**Figure S1d**). At the highest covalent crosslink density (0.05 thiol: monomer ratio) stress relaxation is minimal across all frequencies (**Figure S1d**).

Armed with these design principles, we developed a set of HA hydrogels composed of either HMW (1.5 MDa) or LMW (60 kDa) methacrylated HA exhibiting a range of stress relaxation properties (**Figure 1b**). To benchmark our materials against tissue, we performed stress relaxation measurements on freshly harvested mouse brain (**Figure 1b**, yellow traces), which revealed ∼80% stress relaxation within ∼5 min. While our 30 mg/mL/0.01 thiol: monomer HA hydrogel described above also relaxes stress to ∼80% by 5 min, (**Figure S1d**) it has a much higher elastic modulus than brain tissue (330 vs 200 Pa at 0.5 rad/s). We found that reducing HA concentration to 15 mg/mL while raising the crosslinker concentration from 0.335 to 0.838 mM yielded hydrogels with storage modulus, loss modulus, and stress relaxation closely resembling brain tissue (**Figure 1c**, dark blue traces, hereafter referred to as H80). We supplemented this hydrogel with two comparison formulations. First, we developed a 20% stress-relaxing HMW hydrogel (referred to as H20), in which crosslink density was tuned to set the storage modulus is similar to that of brain tissue at high frequency but relaxes stress ∼4x less than H80 or brain tissue (**Figure 1c**), thus providing the opportunity to detect stress relaxation-dependent changes in cell behavior. Second, as a non-stress-relaxing control, we prepared a covalently crosslinked LMW hydrogel that exhibits minimal (5%) stress relaxation (referred to as L5). We chose LMW HA for this control for two reasons. First, we were unable to find a non-stress-relaxing HMW HA formulation that did not also significantly increase storage modulus. Second, the use of a LMW HA hydrogel serves as a point of comparison with LMW hydrogels commonly used in HA cell culture platforms.^4^ Importantly, the crosslink density of L5 is tuned to yield similar swelling properties to H80 and H20, with similar storage modulus at high frequencies (**Figure 1c**). We subsequently proceeded with H80, H20, and L5 hydrogel formulations to investigate how the degree of stress relaxation in soft hydrogels regulates GBM invasion.

### 2.2 Stress-relaxing hydrogels promote rapid GBM invasion

#### 2.2.1 Qualitative assessment of invasion with GBM spheroid assays

We next applied the two extremes of our HA formulations (H80 and L5) to ask how stress relaxation properties influence 3D GBM invasion. We began by performing tumor spheroid invasion assays (**Figure 2a**) using two GBM culture models: a continuous cell line (U87) and a human patient-derived glioma stem cell (GSC) line (GSC 295). While both cell lines exhibited protrusions (green arrowheads) after culture in L5 hydrogels for 2 days, neither appreciably invaded (**Figure 2b**). By contrast, both U87 and GSC 295 spheroids robustly invaded H80 hydrogels after 2 days, with many single cells emerging from the spheroid (yellow arrowheads) (**Figure 2b**).

**Figure 2.**
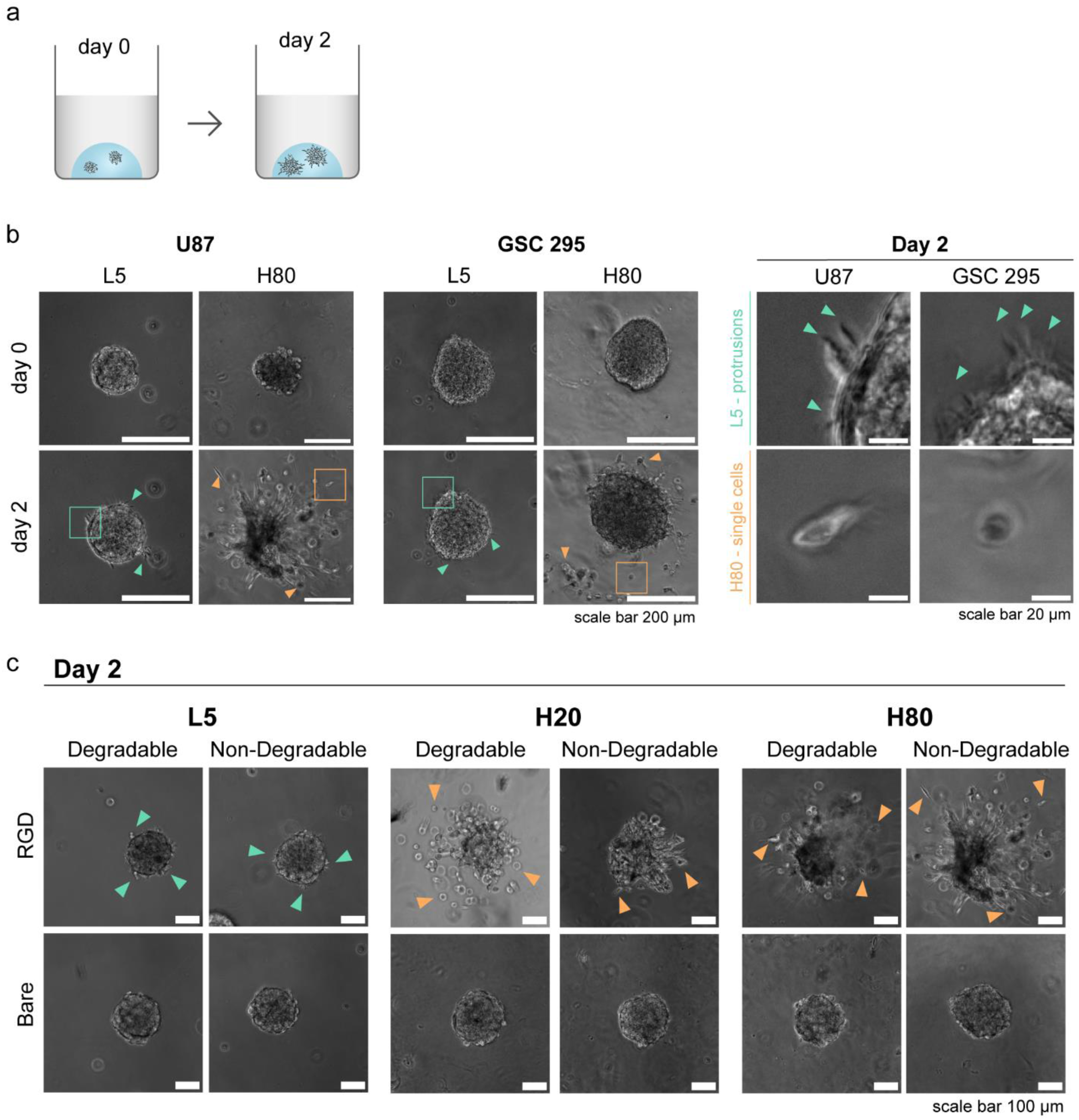
Stress relaxing HMW HA hydrogels promotes rapid invasion. **a.** Spheroid assay schematic. **b.** Representative images of spheroids on day 0 and day 2 from a continuous cell line (U87) and a human patient-derived glioma stem cell (GSC 295) in L5 and H80 hydrogels (scale bar 200 μm). Yellow and green boxes represent inset area. Day 2 insets and arrowheads highlight protrusion-based (green) and single-cell-based (yellow) invasion in L5 and H80 hydrogels, respectively (scale bar 20 μm). **c.** Representative images of U87 cells on day 2 in L5, H20, and H80 hydrogels with either the presence or absence of RGD (RGD/Bare) and the presence or absence of a protease degradable crosslinker (degradable/non-degradable) (scale bar 100 μm). Green arrowheads indicate protrusion-based invasion and yellow arrowheads indicate single-cell-based invasion.

We next asked how stress-relaxing hydrogels influence two biological mechanisms relevant to adhesion and invasion and are commonly tuned within in vitro platforms: RGD-mediated integrin engagement and protease degradation. To address this question, we created 4 variants each of L5, H20, and H80 hydrogels featuring pairwise combinations of the presence or absence of conjugated RGD peptides (RGD vs Bare) and the use of crosslinkers that are either protease-degradable or non-degradable (degradable vs non-degradable). We found that RGD conjugation was critical to rapid invasion in H80 and H20 as well as protrusion formation in L5, whether or not protease-degradable crosslinkers are used (**Figure 2c**). For the remaining experiments in this study, we therefore focused on HA formulations that have both conjugated RGD to support invasion and non-degradable peptide crosslinkers to remove proteolysis contributions to invasion.

*2.2.2 Characterization of rapid invasion in tumoroid devices adapted for confocal imaging* While spheroid-based assays illustrate qualitative differences in invasion across materials and culture models, they suffer from important limitations. For example, the diffuse invasion seen in the most pro-invasive matrices (particularly H80) is challenging to rigorously and reproducibly quantify. Moreover, escape of invasive cells from the imaging field in H80 hydrogels leads to data loss and complicates comparative tracking of invasion across conditions. Thus, we complemented our spheroid assays with studies involving our previously described invasion device in which we seed tumor cells in a channel embedded within a 3D HA hydrogel and allow the cells to invade over several weeks, resulting in a reductionist “tumoroid.”^32,33^ We introduced two device modifications for the current study. First, we affixed the device to a coverslip that can be placed inside a glass bottom dish, thereby facilitating confocal microscopy, and second, we cast the device in a PDMS mold to improve device-to-device reproducibility (**Figure 3a**). The mold additionally enabled us to apply laser cutting to precisely and reproducibly position the cell reservoirs 100-200 μm from the bottom of the coverslip, within the working distance of objectives used in inverted confocal imaging.

**Figure 3.**
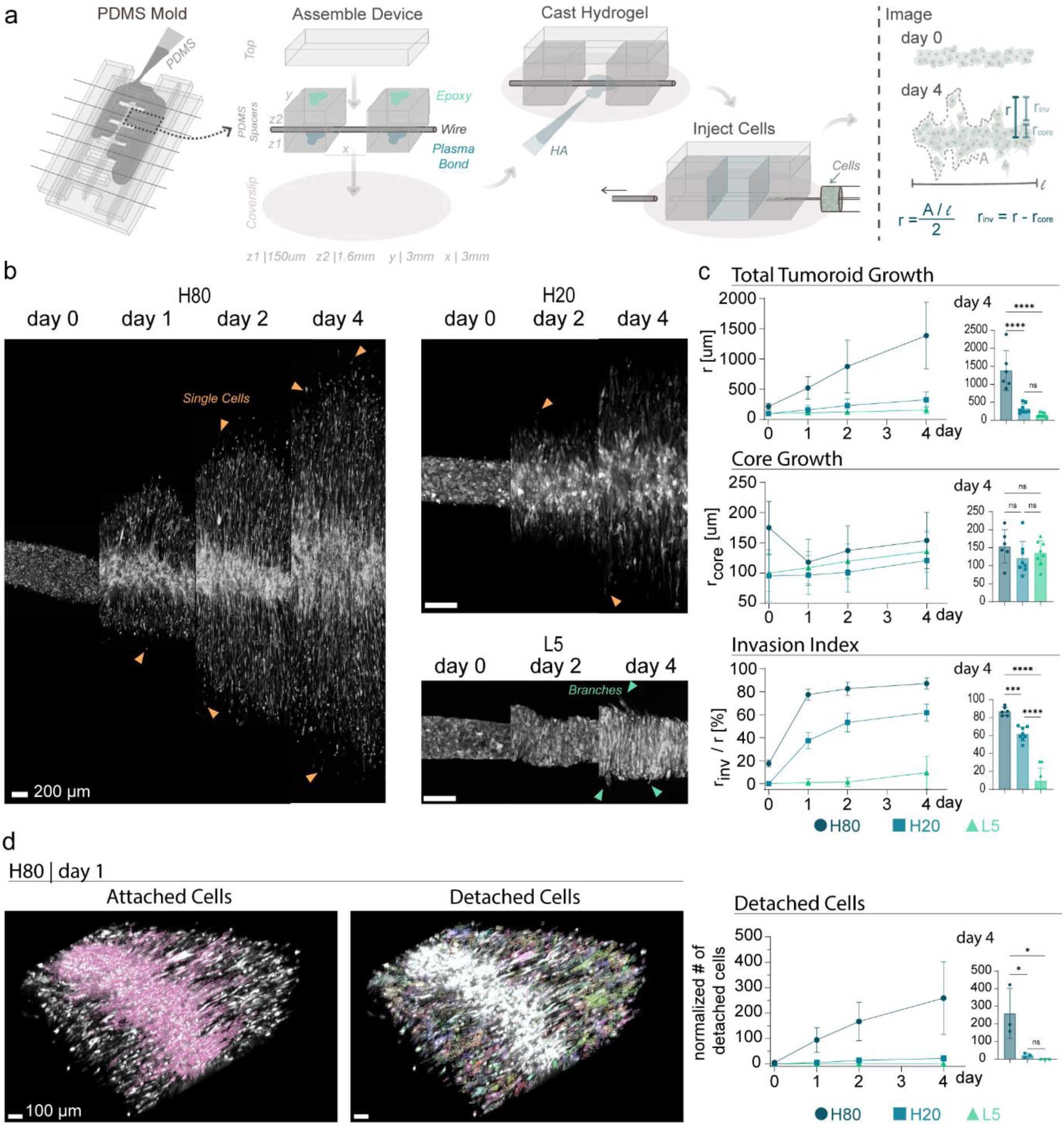
Increasing HA hydrogel stress relaxation leads to faster GBM cell invasion. **a.** Fabrication of the invasion device and schematic of tumoroid images on day 0 and day 4 and variables used to quantify invasion. **b.** Representative z-projections of confocal images of U87 GFP+ cells in the invasion devices from day 0 to 4 (scale bar 200 μm). **c.** Quantification of invasion over time by multiple metrics with statistical analysis at day 4 time point. n = 6-8 devices. ****P<0.0001, ***P<0.001, ns = not significant by a two-tailed one-way analysis of variance (ANOVA) followed by Tukey’s multiple comparisons test. **d.** Quantification of detached cells using Imaris imaging analysis software. Representative images display outlines in different colors around each connected cell cluster (scale bar 100 μm). n = 3 invasion devices. *P<0.05 and ns = not significant by a two-tailed one-way analysis of variance (ANOVA) followed by Tukey’s multiple comparisons test.

We seeded U87 cells expressing a cytosolic green fluorescent protein (U87 GFP+ cells) in these devices and imaged them for 4 days, by which time cells had invaded the full length of H80-based devices. Like the spheroid assay, we observed many single cells (yellow arrowheads) escape the tumoroid core in both H80 and H20 devices (**Figure 3b**). Invasion was also observed in L5 devices, although to a much lesser extent, with cells emerging more collectively from the core, denoted as branches (green arrowheads) (**Figure 3b**). To quantify invasion, we devised a scheme consisting of overall tumoroid radius (r), core radius (r_core_), and invasive radius (r_inv_ = r – r_core_) (**Figure 3a**, Methods). H80 supported much greater overall invasion (r) than H20 and L5, which were statistically indistinguishable from one another (**Figure 3c**). Tumoroid cores in H80 devices initially reduced in volume (decreased r_core_) after 1 day, whereas H20 and L5 tumoroid cores monotonically increased, with all three devices reaching similar core sizes by day 4 (**Figure 3c**). We further quantified invasiveness by calculating an invasion index (r_inv_ / r) and measuring the number of detached cells (**Figure 3c**, **Figure 3d**). Both quantities are greatest in the most stress-relaxing hydrogel, leading us to hypothesize that highly viscoelastic HA promotes rapid invasion by facilitating cell detachment from the core.

### 2.3 Leader cells enable rapid invasion by remodeling stress-relaxing matrices

#### 2.3.1 Leader and follower cell dynamics emerge in the stress-relaxing invasive fraction of H80

To gain mechanistic insight into relationships between stress relaxation, cell detachment from the core, and rapid invasion, we tracked invasion via live-cell confocal imaging of U87 GFP+ cells. Two distinct populations of cells were observed within the invasive fraction, which we term leader and follower cells based on their location and connectivity, analogous to past studies in epithelial systems (**Figure 4a**).^34–37^ Leader cells, which we found at the leading edge of the invasive fraction and are largely unconnected to other cells migrate with a mean speed of ∼25 μm/hr (**Figure 4b**) and are characterized by actin localization at the back of the cell, which could facilitate forward locomotion (**Figure 4c**; white arrowheads).^38–43^ Leader cells also frequently exhibit bead-like protrusions or “pearls,” which appear as early as day 0 as the first set of cells emerge from the core (**Figure 4d**). Past work has shown that pearling protrusions result from the mechanical interplay between the rigidity of the actin cortex and tension produced by a lack of uniform adhesion points.^44–46^ By contrast, follower cells, which trail the leader cells and retain cell-cell contacts exhibit a more elongated morphology with actin-based cables along their lateral borders (**Figure 4c**; white arrowheads), reminiscent of cells migrating in confined channels.^47^ Remarkably, followers can extend protrusions ∼200 μm and ∼4x the length of their cell bodies, extending the leading edge of the cell deeper into the invasive fraction (**Figure 4d**). Process extension is then followed by translocation of the nucleus, which can be discerned as a region of relative GFP depletion (**Figure 4d**).

**Figure 4.**
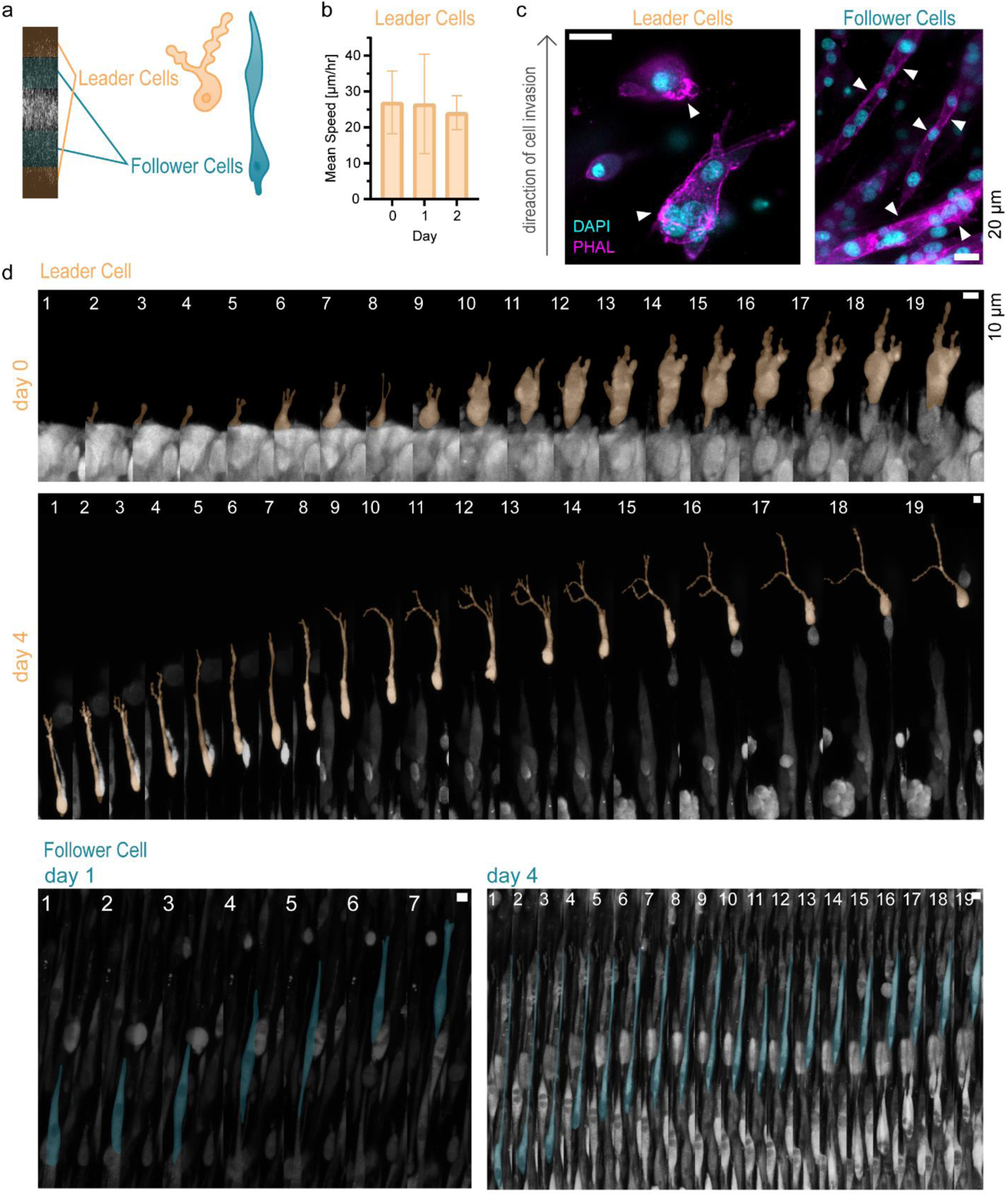
Leader and follower cells in H80 hydrogels. **a.** Schematic of leader and follower cell locations within the invasive fraction. **b.** Mean cell speed of leader cells on days 0, 1, and 2. n = 3 invasion devices, with 3 – 20 cells tracked per device. Means are not significant by a two-tailed one-way analysis of variance (ANOVA) followed by Tukey’s multiple comparisons test. **c.** Phalloidin and DAPI staining of leader and follower cells on day 4 (scale bar 20 μm). Black arrow indicates direction of cell invasion. White arrowheads point out localized actin at the back of the leader cells and along the periphery of follower cells. **d.** Representative montages (frame = 30 min) of confocal z-projections of invading GFP+ leader and follower cells (scale bar 10 μm). Color masks highlight the morphology of cells of interest.

#### 2.3.2 Material properties change within the invasive fraction of H80

The leader-follower invasion dynamic described above led us to explore the hypothesis that leader cells remodel the ECM to facilitate invasion of the followers. To investigate mechanical changes induced by the leader cells, we spatially map cell/ECM viscoelastic properties in the invasive fraction of H80 with atomic force microscopy (AFM). After 4 days in culture, H80 devices were sliced and affixed to a Petri dish with the tumoroid surface face-up, then probed in a series of force measurements along an approximately linear path starting near the tumoroid core and moving radially outward (**Figure 5a, Fig. S2a** points 1-5). The Young’s modulus and degree of stress relaxation depended strongly on the position within the invasive fraction of the tumoroid, with values at the leading edge plateauing at values approaching those of acellular hydrogels (**Figure S2b**). The Young’s modulus was lowest closest to the core, increasing towards the periphery, and plateauing towards the acellular region (∼40-60 Pa) (**Figure 5a**, **Figure S2b**. Conversely, the extent of stress relaxation was highest closest to the core and quickly decreased and plateaued towards the acellular region (25-30% SR) (**Figure 5a, Figure S2**). Young’s modulus did not depend on the proximity of the probe to a given visible cell, indicating that differences in mechanical properties reflect a composite contribution of both cells and ECM (**Figure S2c**). The finding that measurements closest to the core are softer and more stress-relaxing than the periphery or acellular region suggests that cells fluidize the matrix during invasion.

**Figure 5.**
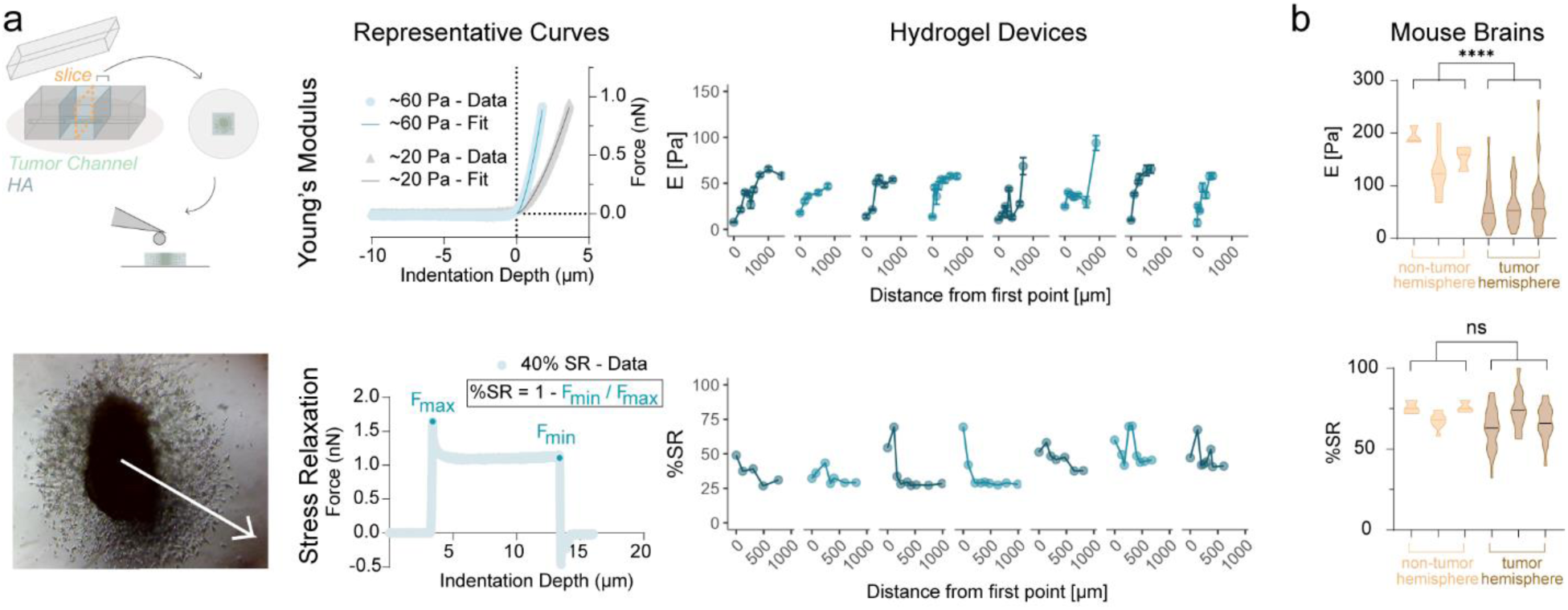
Invading cells remodel the matrix in H80. **a.** AFM measurements of H80 invasion devices on day 4 with a colloidal probe. Schematic and brightfield image display sample preparation and measurement workflow. Representative force curves of Young’s modulus and stress relaxation measurements indicate how AFM force curves were interpreted. Measurements show an increasing trend in Young’s modulus (n = 7 – 16 force curves) and a decreasing trend in stress relaxation (points from single force curves) when measuring from the core radially outward. n = 7-8 paths across 3 devices. **b.** AFM measurements of non-tumor and tumor hemisphere mouse brain slices with a colloidal probe. Each violin bar represents an independent brain slice (n = 3). For each slice, locations were chosen within 5 mm of the tumor core (n = 22-32) or of the contralateral non-tumor hemisphere (n = 3-9). Each Young’s modulus measurement is an average over 7 – 16 force curves and each stress relaxation measurement is from one force curve. ****P<0.0001 and ns = not significant by a nested t-test.

#### 2.3.3 Regional material properties of tumor-laden mouse brain

To compare our in vitro observations to tumor tissue, we implanted mouse SB28-FL GBM cells into the brains of syngeneic mice^48^ and allowed invasive tumors to form over 14 days, then harvested brain tissue and prepared tissue slices for AFM measurements. For comparative purposes, we considered two types of slices: one from the tumor (denoted as tumor hemisphere) and one from the contralateral, grossly tumor-free hemisphere (denoted as non-tumor hemisphere) (**Figure S2d**). After affixing each slice to a Petri dish, we spatially mapped Young’s modulus and stress relaxation throughout each slice and, in the case of the tumor-laden slice, at various distances from the tumor core (**Figure S2d**). As with the devices, the tumor hemisphere was significantly softer than the non-tumor hemisphere (**Figure 5b**). However, at the spatial resolution of the measurement, we did not observe a systematic dependence of either Young’s modulus or extent of stress relaxation on the distance from the tumor core (**Figure S2d**). The extent of stress relaxation did not statistically differ between the tumor and non-tumor hemispheres; however, the tumor hemisphere exhibited a much larger distribution of values (**Figure 5b**), supporting the use of hydrogels with different degrees of stress relaxation to model tumor invasion.

### 2.4 Leader cells mechanically and enzymatically remodel the matrix to facilitate follower invasion

Both our device and brain slice measurements support a model in which cells at the leading edge of the tumor mechanically remodel the matrix to support rapid invasion of follower cells. Our observation that follower cells align in columns and assemble parallelized actin cables further suggests that leader cells may be creating physical channels in the hydrogel through which the followers can migrate. We next sought to more directly explore this possibility.

#### 2.4.1 Stress-relaxing H80 hydrogels do not exhibit significant bulk or microscale viscoplasticity

One mechanism through which leader cells could form channels is microscale plastic deformation of the hydrogel. Rapid invasion had previously been observed in a stress-relaxing IPN of alginate and basement membrane due to the cells plastically deforming a channel of at least 3 μm to allow translocation of the nucleus.^49^ When we conducted bulk creep tests on our HA hydrogels at multiple stresses, we found by contrast that ∼100% of the initial strain was recovered within the H80 hydrogel, whereas only 40 – 70 % of the initial strain was recovered in the brain (**Figure S3a**). These results signify that although the brain exhibits viscoplasticity, the H80 hydrogels are primarily viscoelastic, with little evidence of bulk plastic deformation at this resolution. Seeking evidence of plastic deformation at the cell scale, we obtained live-cell time-lapse images of U87 GFP+ cells in an H80 hydrogel doped with fluorescently tagged HA to track cell-induced HA rearrangements. While HA fluorescence decreases at the rear of the leader cell, it immediately recovers within 30 min, supporting the claim that the matrix is capable of viscoelastic creep, but does not support permanent formation of a channel of up to 3 µm to allow translocation of the nucleus (**Figure S3b**). To improve the resolution of these measurements, we seeded the GFP+ cells in a fluorescent bead-laden H80 hydrogel and tracked bead displacement during invasion. Bead movement was only observed when the bead was in the vicinity of the cell and not observed when the bead was far from the cell (**Figure S3c**), confirming that bead movement is associated with cell motility but not thermal motion. We observed bead motion near the cell to follow a stereotypical cycle: first, as a leader cell invades, the bead is pulled closer to the cell; as the bead and cell come into contact, the bead is pushed away; and finally, once the cell has passed the bead, the bead creeps back to its original position (**Figure S3c**). This sequence of events suggests that while the leader cells do deform the matrix as they invade, the deformation is not entirely plastic or substantial at this resolution.

#### 2.4.3 Leader cells deposit HYAL2-rich trails

Next, we explored the hypothesis that cells are creating minute changes in matrix properties by degrading localized areas in the hydrogel rather than plastically deforming large channels. To do this, we looked more closely at potential degradation-based mechanisms within the invasive front of all the devices. To account for the significantly slower invasion within H20 and L5 at day 4 relative to H80 (**Figure 3c**), we allowed invasion to proceed for 11 days and 18 days in H20 and L5-based devices respectively to match the invasion index of H80 on day 4 (**Figure 6a**). Once the devices reached their endpoints, we stained for hyaluronidase (HYAL) 2, which degrades HMW into LMW fragments and is strongly upregulated in GBM relative to normal brain tissue.^15,50^ We noticed streaks of HYAL2-postivite puncta throughout the invasive fraction, spatially located in tracks between leader and follower cells, in both H80 and H20 (**Figure 6b**). These HYAL2 tracks were also observed in spheroids on day 2 (**Figure S4**). In contrast, HYAL2 was localized to the tips of the invading cells in L5, but since most of the cells are connected to the core with very few detached cells, we did not observe the same HYAL2 trails in L5 as we did in H20 and H80 (**Figure 6b**). Zooming further into the regime between leader and follower cells, we noticed small deposits with brightfield microscopy, which overlap with the HYAL2 trails but not F-actin (**Figure 6c**). The trails seen in brightfield microscopy only present themselves in the wake of a leader cell and remain visible for at least 8 hrs after the leader cell has migrated through the area (**Figure 6d**, top). Since these trails are localized at and behind leader cells, we speculated that leader cells create a trail of deposits with membrane-bound HYAL2 that may promote follower invasion through local HYAL-mediated matrix digestion. The observed brightfield and HYAL2-deposit trails in the devices are very narrow (∼1 μm), consistent with a mechanism in which the leaders provide a directed, column-like path for follower cells to align. We can observe this follower cell alignment in the brightfield cell invasion timelapse, where a follower cell protrusion moves in parallel with the leader cell’s path and by the end of the captured period, more of the cell body becomes visible (**Figure 6d**, bottom), suggesting that the follower cell is elongating and migrating within this path.

**Figure 6.**
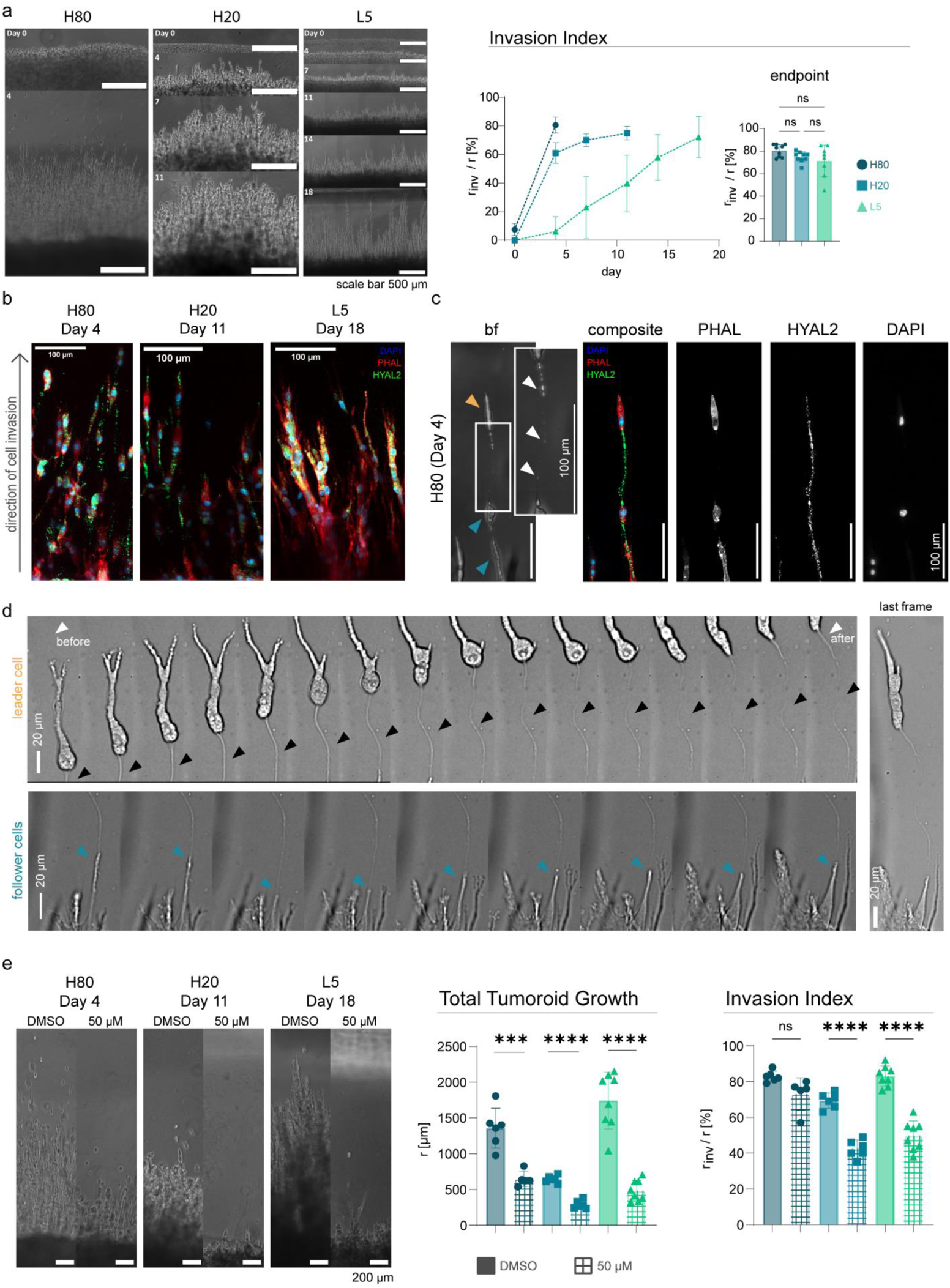
Leader cells prime a path for follower cells invasion. **a.** Representative phase images of invasion devices over time with matched invasion indexes at different endpoints (scale bar 500 μm). n = 8-9 invasion devices. ns = not significant by a two-tailed one-way analysis of variance (ANOVA) followed by Tukey’s multiple comparisons test. **b.** Confocal z-projections of H80, H20, and L5 invasive fractions at their representative endpoints (scale bar 100 μm). **c.** Confocal z-projections and brightfield images of leader-follower dynamics in H80 (scale bar 100 μm). Yellow arrowhead indicates leader cell, blue arrowheads indicate follower cells, and white arrowheads indicate deposits. **d.** Brightfield confocal montages of leader (top, 16 frames) and follower (bottom, last 9 frames) cell invasion and a brightfield confocal image of the final frame (right) on day 4 in H80 (frame = 30 min, scale bar 20 μm, contrast adjustment). As the leader cell invades it leaves a path in its wake; black arrowheads point out the path and white arrowheads show the absence and presence of the path at the beginning and end of the captured time period. Follower cell protrusion aligns itself in the leader cell’s path (blue arrows). **e.** Representative images of H80, H20, and L5 devices at invasion-index endpoints with or without HYAL inhibitor (50 μM or DMSO, respectively), and quantification of total tumoroid growth and invasion index. n = 5-8 invasion devices. ****P<0.0001, ***P<0.001, and ns = not significant by an unpaired two-tailed Student’s t-test.

To determine if HYALs functionally contribute to invasion, we repeated our experiments in the presence of a broad-spectrum HYAL inhibitor (apigenin).^51–53^ Apigenin treatment reduced total tumoroid growth in all the materials, demonstrating the importance of HYALs for cell expansion into the matrix (**Figure 6e**). However, we found that HYAL inhibition did not significantly reduce the invasion index in H80, whereas it did in both H20 and L5 devices (**Figure 6e**), suggesting that the cells employ different modes of invasion in the different hydrogel formulations. Additionally, when we used apigenin in conjunction with H80, follower cells advanced in less well-defined columns, consistent with a weaker leader-followed dynamic (**Figure S5**). This suggests that while invasion in H80 depends much less heavily than HYALs than in H20 and L5, HYALs are important in leader-follower dynamics that support rapid invasion. The combined action of matrix rearrangement and HA degradation gives rise to the leader-follower dynamic we observe in H80. Specifically, we propose that leader cells enter the matrix through a mechanism that involves both mechanical and potentially HYAL-mediated matrix rearrangement. They then leave HYAL2-rich deposits in their wake that open a seam for follower cells to rapidly migrate along. Because H20 and L5 matrices require sustained stress for deformation, the emphasis is shifted towards HYAL-mediated degradation, leading to their slower invasion and bulk expansion (**Figure 7**).

**Figure 7.**
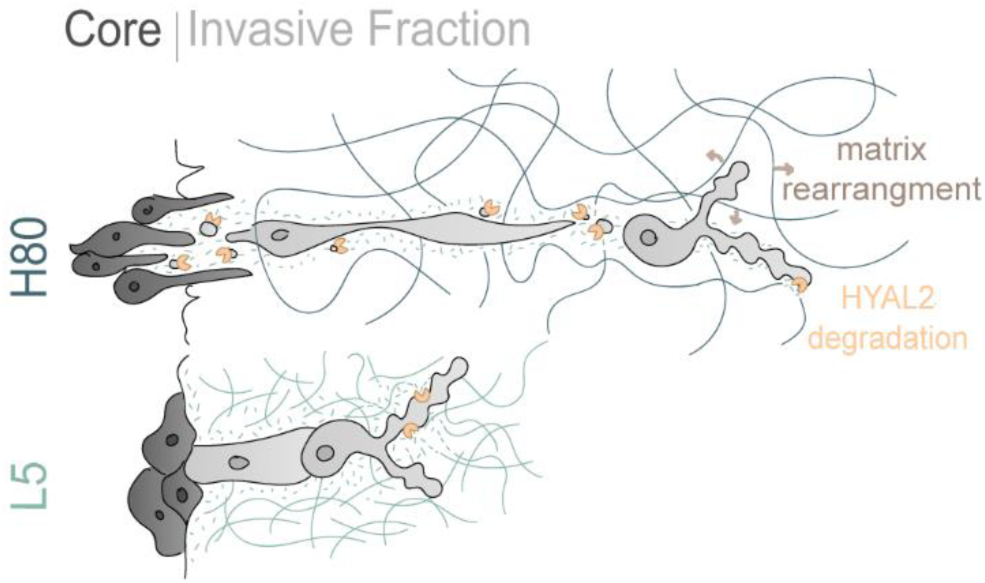
Conclusion. Schematic displaying matrix rearrangement, HYAL-mediated degradation, and leader-follower dynamics in H80 that lead to rapid invasion as compared to L5.

#### 2.4.4 Unbiased microregional RNA sequencing of H80 tumoroids and patient tumors supports a role for HYAL2 and GAG metabolism pathways in invasion

To gain a more complete and unbiased picture of intracellular drivers of invasion in our matrices, we performed mRNA sequencing on invasive and core tumoroid fractions within H80, H20, and L5-based devices.^33^ After the tumoroids reached their respective invasion-index endpoints, they were micro-dissected to separate the core from its invasive fraction based on location and cell density, where the core visibly appeared much darker (**Figure S6a**). mRNA was harvested from each fraction and sent for sequencing. Principal component analysis (PCA) confirmed that all samples within a given group clustered and were more similar to each other than to the other groups (**Figure S6b**).

We focused our analysis on differentially expressed genes (DEGs) in H80 relative to sectioned counterparts in either H20 or L5 (referred to as H80 inv vs H20 inv, H80 core vs H20 core, H80 inv vs L5 inv, and H80 core vs L5 core) (**Figure S6c**). With the four lists of upregulated DEGs from each comparison group, we performed Enrichr pathway analysis using the Reactome (2022) database (**Figure S6d**). Notably, the top pathways for all four comparison groups are associated with ECM metabolism, including cholesterol, glycosaminoglycans (GAGs), steroids, carbohydrates, and fatty acids. While expression of genes relevant to cholesterol metabolism pathways differ between H80 and H20, expression of genes relevant to GAG metabolism differ between H80 and L5. The difference in GAG metabolism is particularly notable given that HA is itself a non-sulfated GAG^15^ and sulfated GAGs are also incorporated on lecticans crosslinking HA within the brain.^15,54–56^ Sulfated GAGs are also incorporated into transmembrane proteins that facilitate adhesion, migration, and binding of growth factors,^57^ which are important in supporting tumor invasion. We observe that many of the GAG metabolism genes, including HYAL2 and HAS3 (an HA synthesis enzyme), are highly expressed in H80 and H20 core and invasive fractions (**Figure S6e**, top), showing the importance of GAG metabolism in viscoelastic, HMW HA hydrogels. Within H80 and H20, we also found many upregulated sulfotransferases (ie HS3ST2A1, CHST14, etc) and sulfatases (SGSH, ARSB, etc), two classes of enzymes that synthesize and degrade GAGs via the conjugation and removal of sulfate groups (**Figure S6e**, top). Some noteworthy upregulated proteoglycans include CD44, an HA adhesion proteoglycan that facilitates invasion, lumican (LUM), which binds collagen molecules within a fibril, and many heparan sulfate proteoglycans (i.e. GPC1, HSPG2, SDC1, AGRN), which are transmembrane proteins that can interact with many ligands (including growth factors, cytokines, and proteinases) (**Figure S6e**, top). GAG metabolism genes are also upregulated in tumor vs non-tumor patient samples, including: LUM, CD44, HYAL2, HSP62, CHST14, HS3ST3A1, GPC1, SDC1, and SGSH (**Figure S6e**, bottom).

## 3. Discussion

HA hydrogels have been increasingly used and proven valuable for modeling biology and disease in the brain. Recent efforts to incorporate viscoelastic effects with dynamic crosslinking into HA hydrogels have lent valuable insight into how viscoelasticity affects cell biology.^25,27,28,58^ However, the range of viscoelastic properties achieved with HA hydrogels has remained somewhat limited and in general not matched the full speed and extent of brain matrix stress-relaxation. In this study we introduce a strategy to fill this gap based on incomplete covalent crosslinking of HMW HA, where the covalent crosslinks provide hydrogel integrity without pure elasticity, and the chain entanglements provide a physical mechanism for stress relaxation. By conducting comparative studies between HMW HA hydrogels with different degrees of stress relaxation (H80 and H20) and LMW HA hydrogels covalently crosslinked to near-complete elasticity (L5), we recapitulate the diversity of stress relaxation properties observed in normal and GBM tumor-laden tissue. We also use this platform to investigate how stress relaxation properties regulated 3D invasion, with the most stress-relaxing formulation (H80) giving rise to rapid “leader-follower” invasion in which leader cells locally digest the matrix with hyaluronidases to create microscale channels that facilitate the rapid invasion of follower cells. This mechanism is supported by fractional transcriptomic analysis, hyaluronidase inhibition studies, and direct imaging. Our work thus both contributes a new material system for modulating stress relaxation properties and lends new insight into how those properties contribute to invasion.

To obtain our most stress-relaxing hydrogel, we leverage polymer MW entanglements as a source of viscous dissipation. Varied MW polymer formulations have previously been used to modulate stress relaxation properties. For example, HMW and LMW alginate hydrogels were applied to investigate effects of stress relaxation on stem cell differentiation.^5^ Interestingly, HMW alginate influenced stress relaxation properties differently than in our system, with the addition of HMW alginate creating entanglements that slowed the rate at which the system can dissipate stress.^5^ The opposite phenomenology may result from the fact that LMW alginate can already stress relax to ∼0 Pa due to its noncovalent Ca^2+^-mediated crosslinks, which can fully dissociate under stress. The entanglements of HMW chains thus function as effective crosslinks that oppose rather than facilitate stress relaxation. For HA specifically, a few studies have explored the effects of HA MW on hydrogel material properties such as stiffness, gelation time, and degradation, but not on viscoelastic properties.^59,60^ Another, another study used semi-interpenetrating networks (sIPNs) of crosslinked gelatin and un-crosslinked HA of varying MW to explore GBM invasion.^22^ While this study did not explore effects of HA MW on viscoelasticity, it did report effects on MW-dependent effects on motility, with faster spheroid growth in sIPNs formed from LMW HAs (10 and 60 kDa) when compared to their HMW counterpart (500 kDa). This result raises the important possibility that HA MW can influence signaling through HA receptors, potentially independent of mechanical contributions of chain length. While the clear differences in invasion between H20 and H80 in our study argue strongly for a role of mechanics in our system, we cannot fully exclude non-mechanical MW effects. Future studies featuring matrices in which HMW concentrations are varied while viscoelastic properties are fixed should help shed light on this issue.

Our system recapitulates invasion patterns previously seen in vivo, such as diffuse invasion, leader-follower dynamics, and dendritic morphologies of leader cells. The fast stress-relaxing nature of H80 allows cells to rearrange the matrix to facilitate the detachment of single cells from the tumoroid core, reminiscent of the diffuse nature of GBM.^16,17^ This detachment not only happens in less than 24 hours, but the resulting plethora of detached leader cells further facilitates rapid invasion through priming a direct path for follower cells. Neither effect is observed to the same extent in L5. This drive to detach from the core can also be observed in the core growth of H80-based tumoroids over time. While the cores of H20 and L5 tumoroids incrementally increase over time, H80 cores immediately decrease before increasing in size, reflecting a “go vs. grow” phenomenon^61^ in which invasion trades off against proliferation. We also observe the invading cells to display morphologies and migration modes reminiscent of those studied in vivo, which have not been robustly recapitulated in past 3D HA hydrogel models. In our material platform, follower cells exhibit similar migration modes as those undergoing translocation^62^ and utilizing the nuclear piston mechanism.^63^ Leader cells also exhibit branching migration and tumor microtube (TM) formation^62^ observed in cells migrating in the brain parenchyma as opposed to the perivascular niche.^64^ We draw parallels in morphology to the TMs seen in vivo to our leader cell protrusions, in that both have irregular morphology and branching.^62,64,65^ The dynamics between the leader and follower cells have also been highlighted in other tumor systems, where MCF10A non-malignant breast epithelial spheroids in viscoelastic alginate hydrogels invade in branched, path-like columns as opposed to their elastic hydrogel counterparts.^66^ These branched, path-like columns are similar to the invasion we observe in our L5 hydrogels as well as the follower cells in our H80 hydrogels. Utilizing the H80 hydrogel platform, we can observe leader and follower morphologies and migration modes relevant to invasion in vivo, which could complement the use of mouse models.

One of our more notable findings is that leader cells leave a trail of HYAL2-rich deposits that prime the matrix for follower cells through local hyaluronidase-mediated digestion. While the precise mechanism through which deposits are released remains unclear, extracellular vesicles (EVs) represent an intriguing possibility,^67–71^ with evidence of EV formation having been observed at the tips of GBM U373 cell protrusions.^72^ EV secretion from protrusions has been proposed to occur during migration, where EV or protrusions potentially stimulate motility through autocrine and/or paracrine signaling.^69^ Indeed, EVs have not only been implicated as a stimulus for directed cell migration,^73^ but also in multicellular migration specifically. Here, EVs are proposed to form at the tips of retracted fibers or cellular protrusions of advancing cells, leaving a trail of EVs in their wake, referred to as ‘footprints’ or ‘adhesive exosome trails’.^74–76^ EVs have also been closely connected with HA-based turnover, adhesion, and signaling; for example, EVs found within spider venom display hyaluronidases, which are capable of digesting HA in solution.^77^ Moreover, some EVs have been shown to be coated with HA, EV secretion increases in HAS3-overexpressing cells, and EVs are enriched in cholesterol (which can bind to and/or modify HA).^67^ Consistent with these observations, our RNAseq data shows enrichment of HAS3 and cholesterol biosynthesis-relevant transcripts in H80. EVs have been implicated in the transport of growth factors between cells to facilitate adhesion and invasion,^67,68^ and in ovarian cancer, matrix-degrading proteinases have been reported to shed from membrane vesicles in vivo and in vitro.^78^ These observations collectively raise the possibility that leader cells in our system secrete EVs or EV-like carriers that promote follower invasion through hyaluronidase-mediated matrix degradation and potentially other paracrine signals. Despite the explosion of interest in EVs in tumor invasion, comparatively little is known of how EV function is regulated by the properties of a 3D matrix.

## 4. Conclusion

While HA hydrogels have been increasingly used to model neuropathological processes, such hydrogels are often composed of covalently bonded LMW HA chains, creating a predominantly elastic system. Recent efforts to incorporate viscoelastic properties into HA hydrogels have led to valuable cell biological discoveries; however, it has proven challenging to formulate a hydrogel that also mimics the fast and full extent of stress relaxation we observe in brain tissue. In our study, we leverage the ability of HMW polymer entanglements to rearrange and reorganize to create and characterize a set of HA-based hydrogels with differing extents of stress relaxation, with the most stress-relaxing hydrogel mimicking the stress relaxation properties of brain tissue. We find that increasing HA stress relaxation leads to faster GBM cell invasion, which occurs through a distinct leader-follower mechanism reminiscent of GBM invasion in vivo. Next-generation sequencing, pharmacologic studies, and microscopy suggest that leader cells use hyaluronidases to enzymatically remodel a narrow path in their wake, facilitating the advancement of follower cells and thus rapid invasion. Our study offers new insight into tuning HA MW to achieve viscoelastic hydrogel properties, demonstrates how these properties drive invasion, as well as argues for the use of highly stress-relaxing materials to model GBM. Our work paves the way to continue to explore relationships between 3D matrix properties, paracrine signaling through EVs and other mechanisms, and rapid leader-follower invasion.

## 5. Methods

### 5.1. Tissue Culture

U87 MG human GBM cells were obtained from the University of California, Berkeley Tissue Culture Facility, which sources its cultures directly from the American Type Culture Collection. U87 cells were cultured in fully supplemented high-glucose Dulbecco’s modified Eagle medium (DMEM, Gibco) supplemented with 10% (vol/vol) fetal bovine serum (Corning), 1% (vol/vol) penicillin-streptomycin (Gibco), 1% (vol/vol) sodium pyruvate (Gibco), and 1% (vol/vol) MEM non-essential ammino acids solution (Gibco). U87 cells were transfected with cytosolic green fluorescent protein (GFP) for easy visualization. Bevacizumab-sensitive/resistant U87 cells were cultured identically to wild-type U87 cells.^79^ Cells were screened for mycoplasma every 3 – 4 months with the MycoSensor qPCR Assay Kit (Agilent Technologies).

GSC-295 cells were obtained from The University of Texas M.D. Anderson Department of Neurosurgery. GSC-295 cells were propagated as neurospheres in DMEM/F12 basal medium supplemented with 2% (vol/vol) B-27 supplement (Gibco), 20 ng/mL EGF (R&D Systems), and 20 ng/mL FGF (R&D Systems). For experiments on HA gels, medium was supplemented with 0.1% penicillin/streptomycin (Gibco).

Murine glioblastoma SB28-FL were generously provided by Dr. Hideho Okada, University of California San Francisco, San Francisco, CA. SB28-FL cells were cultured in RPMI 1640 medium supplemented with 10% (vol/vol) fetal bovine serum, 2% (vol/vol) GlutaMAX (Gibco), 1% (vol/vol) non-essential amino acids (Gibco), 1% (vol/vol) hydroxyethyl piperazineethanesulfonic acid (Gibco), and 1% (vol/vol) penicillin-streptomycin (Gibco).

### 5.2. Animal studies

For SB28 in vivo studies, eight-week old C57BL/6 mice were purchased from the Jackson Laboratory. Animals were housed in UCSF under pathogen-free conditions, and all in vivo protocols were approved by UCSF IACUC (AN201734-00B). Ten thousand (10,000) SB28-FL tumor cells in two microliters were implanted intracranially using a stereotactic frame. The following coordinates were used from the bregma: anteroposterior (AP), 0 mm; mediolateral (ML), 1.9 mm; and dorso-ventral (DV), 3.0 mm. Bioluminescent imaging (Xenogen IVIS Spectrum) was conducted on day seven to verify tumor engraftment and characterize tumor size. Mice were euthanized on day fourteen, and whole brain tissue was collected for AFM measurements.

### 5.3. HAMe synthesis

HA polymers were methacrylated as previously described.^4^ Briefly, methacrylic anhydride (Sigma-Aldrich, 94%) was used to functionalize sodium hyaluronate (Lifecore Biomedical, Research Grade; LMW 66 kDa-99 kDa and HMW 1.5 MDa) with methacrylate groups, referred to as HA-methacrylate (HAMe). A few modifications to the original protocol were made for HMW HA methacrylation and its use in cell culture. Due to the high viscosity of HMW solutions, the methacrylation reaction was carried out in a larger round bottom flask (e.g. 1 L flask for 1 g HA), thus enabling the unmodified polymer powder to be diluted in DI water to 2-3 mg/mL overnight. Conversely, LMW HA can be suspended at 5-10 mg/mL immediately prior to the reaction (e.g. 0.5L flask for 1 g HA). When adding NaOH to buffer the reaction to a pH between 8 – 9, drops were added at smaller volumes and more frequently in HMW than LMW reactions to allow adequate time for mixing without overly alkalinizing the solution and risking backbone hydrolysis. In the purification steps after precipitating HAMe with EtOH, HMW takes longer to redissolve, which may be sped by breaking up the precipitated solids into smaller chunks, applying heat, and mixing in a round bottom flask. Because HMW HA solutions clog filters normally used for vacuum filtration, we centrifuge at 4000 x g for 1 hr to spin down unwanted particles. Lastly, when using HMW solutions in cell culture, it is essential to use concentrations 40 mg/mL or lower to avoid gelation within the tube and to use positive displacement pipettes. All HMW solutions should be pipette mixed rather than vortexed, as well as centrifuged to remove bubbles.

The extent of methacrylation per disaccharide was quantified by ^1^H NMR as detailed previously and found to be 80-100% for materials used in this study. To add integrin-adhesive functionality, HAMe was conjugated via Michael Addition with the cysteine-containing RGD peptide Ac-GCGYGRGDSPG-NH2 (Anaspec, AS-62349) to obtain HAMe-RGD stock solutions used to make 15 mg/mL HA hydrogels that have a final RGD concentration of 0.15 mg/mL.

### 5.4. Hydrogel crosslinking

To form 15 mg/mL HA hydrogels, HAMe or HAMe-RGD were crosslinked in phenol-free serum-free DMEM (Gibco) with di-thiol peptide crosslinkers that were either broadly protease-degradable (KKCGGPQGIWGQGCKK, Genscript) or non-degradable (KKCGGDQGIAGFGCKK, Genscript). HA hydrogels had a total peptide crosslinker concentration of 0.838 mM for H80, 1.676 mM for H20, and 4.610 mM for L5. For protease-degradable hydrogel conditions, H80, H20, and L5 all had a 0.838 mM concentration of the degradable crosslinker, with the non-degradable crosslinker providing the remaining concentration for H20 and L5.

Fluorescent markers were embedded in H80 hydrogels to visualize matrix remodeling. To visualize HA, methacrylated HA was functionalized with cysteine-TMR (Genscript) at a concentration of 0.075 mM. Bead-laden hydrogels were used to visualize matrix creep and plasticity, where sulfate-modified (580/605) 1 μm sized FluoSpheres^TM^ (Thermo Fisher, F8851) were mixed into the hydrogel solution prior to crosslinking, for a final concentration of 0.05% solid.

### 5.5. Material characterization

The shear moduli of hydrogel formulations were measured using a stress-controlled oscillatory rheometer (Anton Parr Physica MCR 310). Briefly, 0.5 mm thick hydrogels were crosslinked for 2 hr in a humidified 37°C chamber. After gelation, hydrogels were cut and placed between an 8 mm parallel plate and lowered to a gap height of ∼0.5 mm. Samples were enclosed in a humidified 37°C chamber. Rheological testing consisted of frequency sweeps ranging from 50 to 0.5 rad/s at 0.5% amplitude. Stress relaxation tests were conducted at 15% strain for 5 min, measuring stress every 0.5 s. All moduli are reported as the average of at least 3 tests per matrix composition.

Swelling tests were conducted by swelling a 100 μL hydrogel in a non-TC treated 35 mm Petri dish in media for 3 days in a humidified 37°C chamber. The swollen hydrogels were measured (m_swollen_) and then washed with DI water until no media color was observed. They were then frozen, lyophilized, and then re-measured (m_dry_) to obtain the swelling ratio, Q: (m_swollen_-m_dry_) / m_dry_.

### 5.6. Spheroid assay

Tumor spheres were assembled using AggreWell Microwell Plates (Stemcell Technologies). Briefly, 2.4×10^5^ cells were seeded into a single well of the Aggrewell plate with 2 mL of media per well to form spheroids consisting of 200 cells. After a 96-hour incubation, spheroids were resuspended in media and mixed with the HAMe, DMEM, and crosslinker to give ∼2-5 spheroids per hydrogel. For phase contrast imaging, 10 μL hydrogel droplets were plated in a 24 well non-TC treated plates, and for immunostaining, 3 uL hydrogel droplets were plated in a 96 well glass bottom plate. Plates were then flipped to maintain the spheroids in the middle of the hydrogel while crosslinking for 2 hrs in a humidified 37°C chamber. After gelation, media was added to the wells and phase contrast images were captured using an Eclipse TE2000-E Nikon Microscope with a Plan Fluor Ph1 10x objective on day 0 and day 2.

### 5.7. Invasion devices

Devices were fabricated following a modified version of our previously published invasion device protocol.^32^ The invasion devices are comprised of a 12 mm No.1 cover glass (Fisherbrand) as the base, PDMS spacers on the sides with a wire through the middle, and a laser-cut acrylic lid on top. To fabricate the lid and the mold for the PDMS spacers, acrylic pieces are laser-cut out of 1.5 mm thick CLAREX° acrylic glass (Astra Products). The PDMS spacer mold is held together using double sided tape (3M) and binder clips. Once 0.00695 mm outer diameter cleaning wires (Hamilton, 18302) are inserted into the mold, Polydimethylsiloxane (PDMS) is poured into the mold and cured for at least 4hr in an 80°C oven. PDMS was fabricated by mixing a 10:1 mass to mass ratio of Sylgard 184 elastomer with the initiator (Dow Corning). PDMS spacers are removed from the mold, sliced in half, and plasma bonded to the cover glass. A slightly smaller 0.00695 mm outer diameter cleaning wire (Hamilton, 18300) is secured inside the spacers to serve as the channel mold to provide a smaller cell reservoir for confocal imaging. The lid is epoxied to the PDMS and let to cure overnight. The devices are then UV-treated for 30 min and stored in cold room prior to use.

The day before the experiment, devices were brought to room temperature and HA hydrogels were casted around the wire and incubated for 2 hrs in a humidified 37°C chamber. After crosslinking, devices were submerged in phenol-free serum-free DMEM (Gibco) overnight. The next day, the wires were removed from the devices and 10,000 – 30,000 cells were seeded into the open channel and the channel ends were plugged with a wire. Devices were then vacuum greased to the bottom of a well plate and bathed in 1 mL media. Devices were cultured for either 4 days or until their respective invasion-index endpoint, and media was changed every 2-3 days.

Invasion was tracked over the course of 4 days or until the invasion index endpoint. Phase contrast images were captured using an Eclipse TE2000-E or Eclipse Ti2 Nikon Microscope. Large images were obtained using a 6 x 6 matrix of a Plan Fluor Ph1 10x objective with a 10% overlap. Confocal images were captured using a Zeiss Cell Discoverer 7 microscope with Airyscan2 technology and a Plan-Apochromat 10x/0.35 objective. Z-projections were obtained using orthogonal projection on the Zeiss Zen Blue software processing with the standard deviation setting. Confocal brightfield images were adjusted in ImageJ by enhancing contrast.

### 5.8. Invasion quantification

In ImageJ, total tumoroid and core areas (A) were outlined on z-projections for each device at each timepoint and channel lengths were measured (l). Core areas were differentiated based on location and cell density, where the intensity of the core visibly appeared much brighter. Radi were calculated by the following equation: r = A / (2 * l), where r = radius, A = area, and l = length of the channel. Radius of the invasive fraction was then calculated by the following equation: r_inv_ = r – r_core_., where r_inv_ = radius of the invasive fraction, r = radius of the total tumoroid, and r_core_ = radius of the core. The invasion index was calculated using the following equation: r_inv_/r.

Imaris imaging software was used to analyze the number of detached cells and leader cell speed. For each 3D image, surfaces were created, where thresholding and background subtraction were used to isolate individual objects. Volumes less than 900 μm^3^ were filtered out. The number of individual object IDs other than the core were recorded as the number of detached cells or cell clusters. Number of detached cells were normalized to a channel length of 500 μm and a channel height of 100 μm. For each 4D image, 10 μm points were manually created in the center of each leader cell. Points from the same cell across multiple timepoints were combined to create a track, and statistics of cell tracks were exported for plotting. Velocity of each frame was calculated as the delta displacement length divided by the time between frames, which was then averaged among all the time frames for each cell. Within a device, mean speed was calculated as the average among all the leader cells, with a minimum of 3 leader cells, and then mean speed for each day was calculated as the average among all the devices.

### 5.9. AFM measurements

All samples were sliced, affixed to Petri dishes, and measured under room-temperature phosphate-buffered saline (PBS) within 2 hr of slicing (hydrogel devices) or 8 hr of animal sacrifice (brains). Devices were grown for 4 days in a modified version of the invasion device, where the hydrogel casting area increased to 8 mm x 8 mm x 8 mm to allow for easier device handling during slicing and adherence steps. After 4 days in culture, the devices were disassembled, sliced, and affixed to a petri dish using Poly-D-lysine (Sigma-Aldrich). For each brain, 4 slices were acquired from 3 cuts: the first was straight down the center of the incision mark located in the center of the right cortex, the second was down the middle of the brain separating the left and right hemisphere, and the last was on the side contralateral to the tumor (left hemisphere). Slices were affixed to the petri dishes using 0.1% agarose (Fisher Scientific).

Measurements were performed with an MFP-3D-BIO AFM (Oxford Instruments) mounted on an inverted optical microscope (Nikon Eclipse Ti) using colloidal probes. Colloidal probes were made according to published protocols.^80^ Briefly, tipless cantilevers (PNP-TR-AU-TL, NanoWorld, nominal spring constant 0.08 N/m) were calibrated using the thermal method and one 25-μm diameter polystyrene sphere (07313-5, Polysciences) was affixed to the end of each cantilever with a small amount of UV-cured glue (AA 349, Loctite).

Young’s modulus measurements on hydrogel devices used a trigger force of 1 nN and force distances of 5-20 μm depending on sample adhesion, to ensure detachment before the next measurement. Ramp rate was changed accordingly to maintain a constant tip velocity of 4 μm/s for all measurements. At each location, replicate force curves were performed in a 4×4 or 8×8 force map with measurements spaced by 1 μm in both X and Y, and a corresponding image was taken with the optical microscope (10x objective) for location tracking. For hydrogel device stress relaxation measurements, single measurements were taken at each location along with an optical image. Using the indentation ramp feature of the AFM manufacturer’s software, the AFM was programmed to approach the sample at a velocity of 4 μm/s until a trigger force of 0.1-0.15 nN, then to indent the sample at a fast rate of 20 μm/s to 3 μm indentation depth (calculated real-time as Z signal minus deflection), and to maintain that indentation for 10 s while measuring deflection before retracting from the surface. The rate of 20 μm/s was determined to be the fastest rate possible without overshooting the indentation depth. For brain samples, Young’s modulus and stress relaxation measurements were taken in the same manner but with a trigger point of 3 nN for Young’s modulus measurements. Tip velocity was maintained at 4 μm/s.

Raw data files were imported to MATLAB and read with a user generated package.^81^ After converting Z piezo position to separation distance and setting the pre-contact baseline force to 0 nN, contact point was determined using the method of Domke and Radmacher.^82^ Force curves were fitted to a Hertzian model for a spherical probe to extract Young’s modulus:^83^

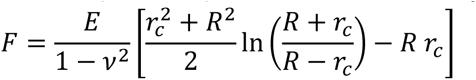

Where 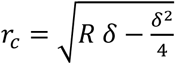is the contact radius, δ is the indentation depth relative to the contact point, R is the spherical particle radius, E is the sample Young’s modulus, F is the measured force, and ν is the sample Poisson ratio which is assumed to be 0.5.

For stress relaxation measurements, extent of stress relaxation was defined as 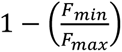, where Fmin and Fmax are the minimum and maximum measured force values within the 10 second relaxation time.

For hydrogel device AFM location data, relative distances were determined with ImageJ for invasion paths (**Fig. S2b**, points 1-5) and XY locations were determined with Adobe Illustrator for tumorous brain slices (**Fig. S2d**). For hydrogel invasion paths, tip location images were stitched together on ImageJ and the shortest distance between each point and the first point was measured, where the first point occurred as close to the core center as a measurement could be taken. Brain slices were affixed to a Petri dish marked with a large “x” in the center that spanned the size of the dish, and then imaged on a printed 0.5 mm grid. The image was uploaded to Illustrator, where a 0.5 mm grid was overlayed on top of the image. 0.5 mm rulers were placed on the AFM X and Y tracks, so all movements would be in increments as low as 0.25 mm up/down and left/right. The “x” was located on the optical microscope and its X and Y coordinates on the rulers were then recorded on Illustrator to establish the spatial location of the “x” crossover point on the tumorous slice. Subsequent measurements followed this XY location approach in order to spatially map points to locations on the tumorous slice.

### 5.10. Immunostaining

Prior to immunostaining, invasion devices and spheroids were fixed for 15 minutes in 4% paraformaldehyde (PFA, Alfa Aesar) with agitation and rinsed three times with 1X PBS and stored in the cold room until staining. Samples were thawed for 10 min at room temperature, permeabilized with 0.05% Triton X-100 (Millipore Sigma) and blocked with 5% Goat Serum (GS, Sigma-Aldrich) in PBS for 1 hour at room temperature. Primary antibodies were added at 1:200 dilution in 1% (vol/vol) GS in PBS and incubated for 3 days. Secondary antibodies were added at 1:500 dilution in 1% (vol/vol) GS in PBS with DAPI and phalloidin stains, following manufacturer’s protocol, and incubated overnight. Both incubations occurred in the cold room with agitation and protected from light. Three PBS rinses occurred after each step. Primary antibodies used were rabbit polyclonal anti-HYAL2 (Abcam, ab68608). Secondary antibodies used were goat anti-rabbit secondary Alexa Fluor 488 antibody (Thermo Fisher Scientific, R37116). Stains used were DAPI (Sigma-Aldrich, 10236276001), Alexa Fluor 546 phalloidin (Thermo Fisher Scientific, A22283).

### 5.11. HYAL inhibitor study

Small molecule HYAL inhibitor Apigenin (Selleck Chemicals) was reconstituted and handled according to manufacturer recommendations. For 3D cell viability assays, cells were seeded at 20k cells/mL in 3 μL gels in a 96 well glass bottom plate and incubated in a humidified 37°C chamber for 2 hrs while crosslinking. After crosslinking, hydrogels were submerged in media containing Apigenin at concentrations 0.1 – 500 μM. After a 48-hour incubation, a LIVE/DEAD Cell Imaging Kit (ThermoFisher) was used to fluoresce live cells (ex/em 488 nm/515 nm) with calcein AM and dead cells (ex/em 570 nm/602 nm) with ethidium homodimer-1. Z-stacks were captured for each hydrogel (x3 gels / concentration) on a Nikon Eclipse Ti2 epifluorescence microscope. Using ImageJ, 3 random areas were selected within each z-stack to create a z-projection and threshold. The number of live cells (cells_live_) and number of dead cells (cells_dead_) were counted for each image, and cell viability was accessed via the following formula: cells_live_ / (cells_live_ + cells_dead_). 50 μM was chosen for future experiments as the highest dose that didn’t affect cell viability compared to the control (**Figure S7**). Devices were cultured in media with either 50 μM Apigenin or a DMSO control, and media was replenished every 2 days. Phase contrast images were captured using an Eclipse Ti2 Nikon Microscope on day 0 and at the invasion index endpoint. Large images were obtained using a 6 x 6 matrix of a Plan Fluor Ph1 10x objective with a 10% overlap, which were analyzed using the same invasion index and total tumoroid growth method stated above.

### 5.12. RNA extraction

For RNA extraction of cells in invasion devices, devices were carefully disassembled using a razor blade and the invasive cells were physically separated from the non-invasive “core” cells using a scalpel and tweezers. All samples were placed in Eppendorf tubes and treated with 10k U/ml Hyaluronidase from bovine testes, Type IV-S (HYAL4, Sigma-Aldrich) for 30 min with agitation at 37°C until the hydrogel was fully degraded and pipette-able. Samples were spun down and excess media containing degraded HA and HYAL4 was removed, leaving a cell pellet. Cells were lysed using 100uL of TRIzol Reagent (Invitrogen), agitated via pipetting and vortexing for 30 seconds, and then freeze-thawed in the-80°C overnight. RNA was then extracted and purified from the cell lysates using Direct-zol RNA MicroPrep (Zymo) following the manufacturer’s protocol with DNase treatment. RNA was dissolved in RNase-free DNase-free distilled H2O and RNA concentration and purity was measured with the Nanodrop. RNA sequencing samples were frozen in the-80°C and sent on dry-ice to Novogene.

### 5.13. RNA sequencing analysis

Isolated RNA was sent to Novogene Corporation Inc. (Sacramento, CA) for library construction, quality control, sequencing, and data filtering. Statistically significant upregulated differentially expressed genes (DEGs) were identified using DESeq2 and chosen based on a statistical cutoff of |Log2FC| > 0 and p-value < 0.05. We used Enrichr^84–86^ to perform pathway enrichment analysis of DEG lists using the 2022 Reactome^87^ pathway databases. Pathways were ranked by their average overlap ratio (DEGs in the pathway: total number of genes in the pathway), giving favor to the pathway that has more dots for the same overlap ratio value. Pathways are chosen based on the top 4 pathways for each DEG list based on adjusted p-value (only adjusted p-values < 0.05 are displayed). Dot sizes are chosen based on their adjusted p-values within each list (ranked 1 – 4 with 1 being the smallest adjusted p-value value and largest dot size). Log_2_ fold change of tumor vs non-tumor genes from the Rembrandt^88^ data set (obtained from the GlioVis data portal (http://gliovis.bioinfo.cnio.es/) was calculated.

### 5.14. Statistics

Statistical analyses of the data from this study were generated using GraphPad Prism 10. The data is presented as the means with error bars for standard deviation. Either an unpaired two-tailed Student’s t-test, or nested t-test was used for comparison between two groups. For multiple comparisons, two-tailed one-way analysis of variance (ANOVA) followed by Tukey’s multiple comparisons test was conducted. Throughout the study, each condition was performed with at least 3 samples, and one to three independent experiments were conducted to ensure reproducibility of the results. At least 3 biological replicates were conducted for each condition. For all statistical tests unless otherwise noted: *P < 0.05, **P < 0.01, ***P < 0.001, ****P < 0.0001.

## Acknowledgements

Confocal images were acquired at the CRL Molecular Imaging Center, RRID:SCR_017852, at the University of California, Berkeley (UC Berkeley), which provided the Zeiss Cell Discoverer 7 microscope with Airyscan2 technology and the Imaris imaging software. We thank Holly Aaron and Feather Ives for their microscopy advice and support. Additional Imaris support was supported by the UC Berkeley Biological Imaging Facility. We thank Dr. Denise Schichnes for her assistance and training. Glioma Stem Cell lines were developed at the University of Texas M. D. Anderson Department of Neurosurgery support by grants from the National Cancer Institute (1R01CA214749, 1R01CA247970, P30CA016672 and 2P50CA127001) and the University of Texas M. D. Anderson Moon Shots Program^TM^. We also thank the following groups and individuals: Garrett Dempsey for providing and dissecting mouse brain samples for stress relaxation and oscillatory rheology; Dr. David Schaffer, Dr. Hyuncheol Lee, and Ana Carneiro for providing and dissecting mouse brain tissue samples for creep tests; Erin Akins for providing the U87 cells expressing cytosolic GFP; Kwasi Amofa for culturing the GSC-295 spheroids; and Katherine Patterson for mycoplasma testing. Finally, we gratefully acknowledge financial support from the following sources: NIH Grants R01CA260443 and R01GM122375 (to S.K.), R01CA227136 (to M.K.A. and S.K.), R01118940 (to S.K. and A.S.); and NSF GRFP (to E.M.C).

## Data Availability

Data shown for this paper is available upon request.

## Conflict of Interest Statement

The authors declare no competing interests.

Received: ((will be filled in by the editorial staff))

Revised: ((will be filled in by the editorial staff))

Published online: ((will be filled in by the editorial staff))

## Supporting Information

### Supplemental Methods

#### Creep measurements

70 uL hydrogels were crosslinked on the rheometer using a 25 mm diameter 1° angle cone and plate probe and with a hydration chamber. For the oscillatory amplitude sweep, after hydrogels were crosslinked, they were subjected to a series of increased strains between 0.1 – 100 % strain, and then resulting stress was measured. For creep tests, after hydrogels were crosslinked, they were exposed to 0 Pa stress for 1 min to observe a baseline strain, then subjected to a step stress of either 10, 30, 50, or 150 Pa for 20 min, and then stress was removed and strain was recorded for 10 min.

For mouse brain samples, ∼0.5 – 1 mm thick slices were cut using a razor blade, and then 8 mm disks were created using a biopsy punch. Samples were placed on the rheometer, and an 8 mm parallel plate probe was lowered until it made contact with the brain slice to create a normal force slightly > 0.0 N. Samples were first subjected to a 0.5% strain at 1 rad/s for 1 min and then exposed to 0 Pa stress for 1 min to create a baseline strain. For the oscillatory amplitude sweep, samples were subjected to a series of increased strains between 0.1 – 100 % strain, and then resulting stress was measured. For creep tests, samples were subjected to a step stress that was held for either 1 min or 10 min and then removed for either 0.5 min or 5 min. Step stresses implemented were 2 and 10 Pa for the 1 min hold and 5 and 30 Pa for the 10 min hold.

#### Mouse brain samples for rheology

Brain tissue samples collected for oscillation and stress-relaxation measurements were from ∼7.5 month old mice, breed (C57/bl6 from JAX: The Jackson Laboratory, all male). Brain tissue samples collected for creep measurements were from ∼2 year old mice, breed (C57BL/6-DhhCre, mix of male and female). Following dissection, tissue samples were transferred to HEPES buffered media and placed on ice. HEPES buffered media contained XF base media (Agilent 103334-100) supplemented with 10% FBS, 25 mM glucose, 2 mM glutamine, 1% Pen/Strep, and 25 mM HEPES for a final pH of 7.4. Samples were cut with a razor blade to create ∼1 mm thick slices, and then punched out with an 8 mm biopsy punch.

**Figure S1.**
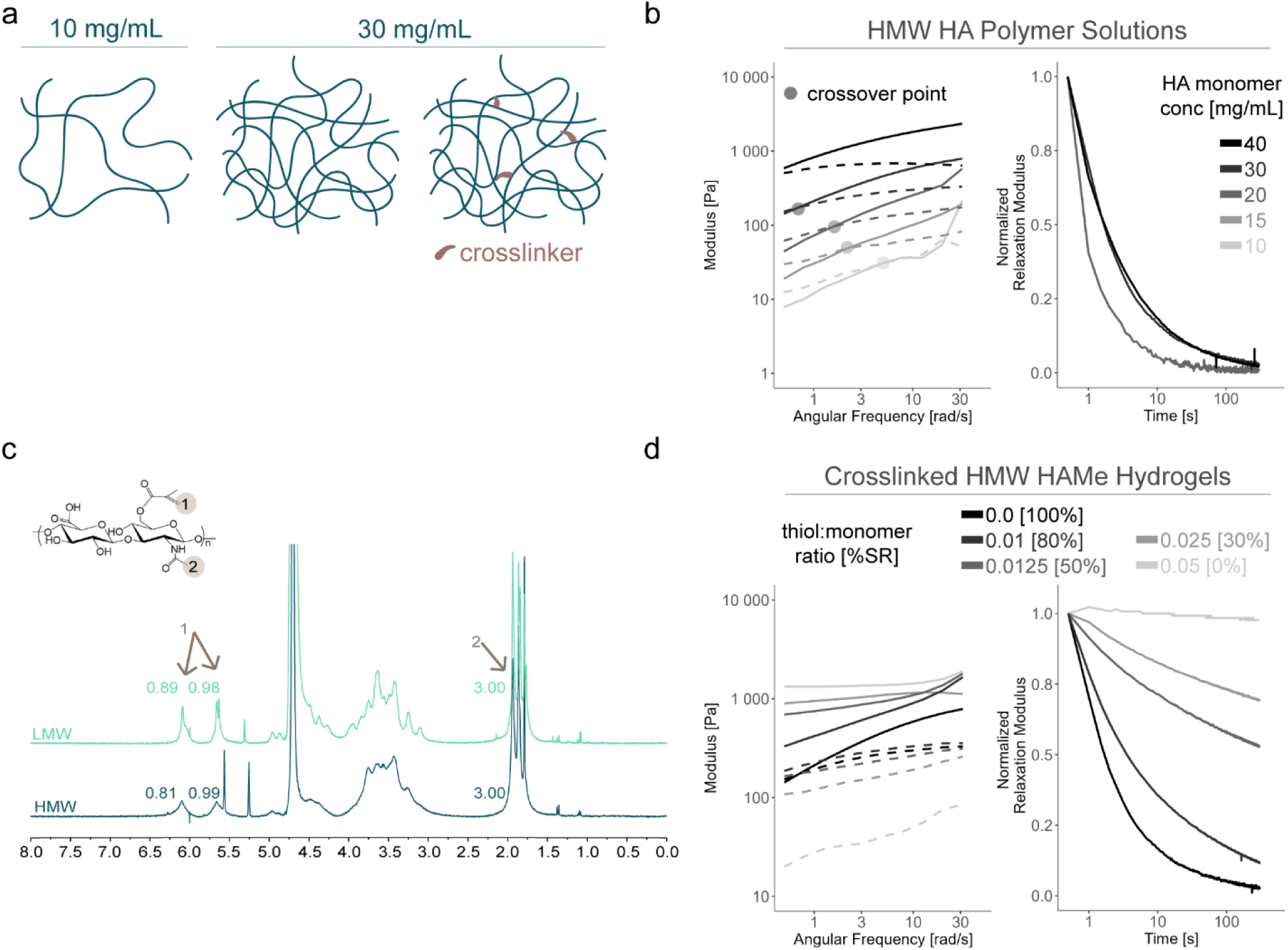
HMW HA entanglements. **a** Schematic of HMW HA polymer solutions at two different HA monomer concentrations, with and without covalent crosslinks. **b.** Oscillatory and stress relaxation tests for HMW polymer solutions at different HA monomer concentrations. **c.** NMR of methacrylated LMW and HMW polymers show high (80 – 100%) methacrylation for both polymers. **d.** Oscillatory and stress relaxation tests for 30 mg/mL HMW polymer mixtures with varying amounts of covalent crosslinker.

**Figure S2.**
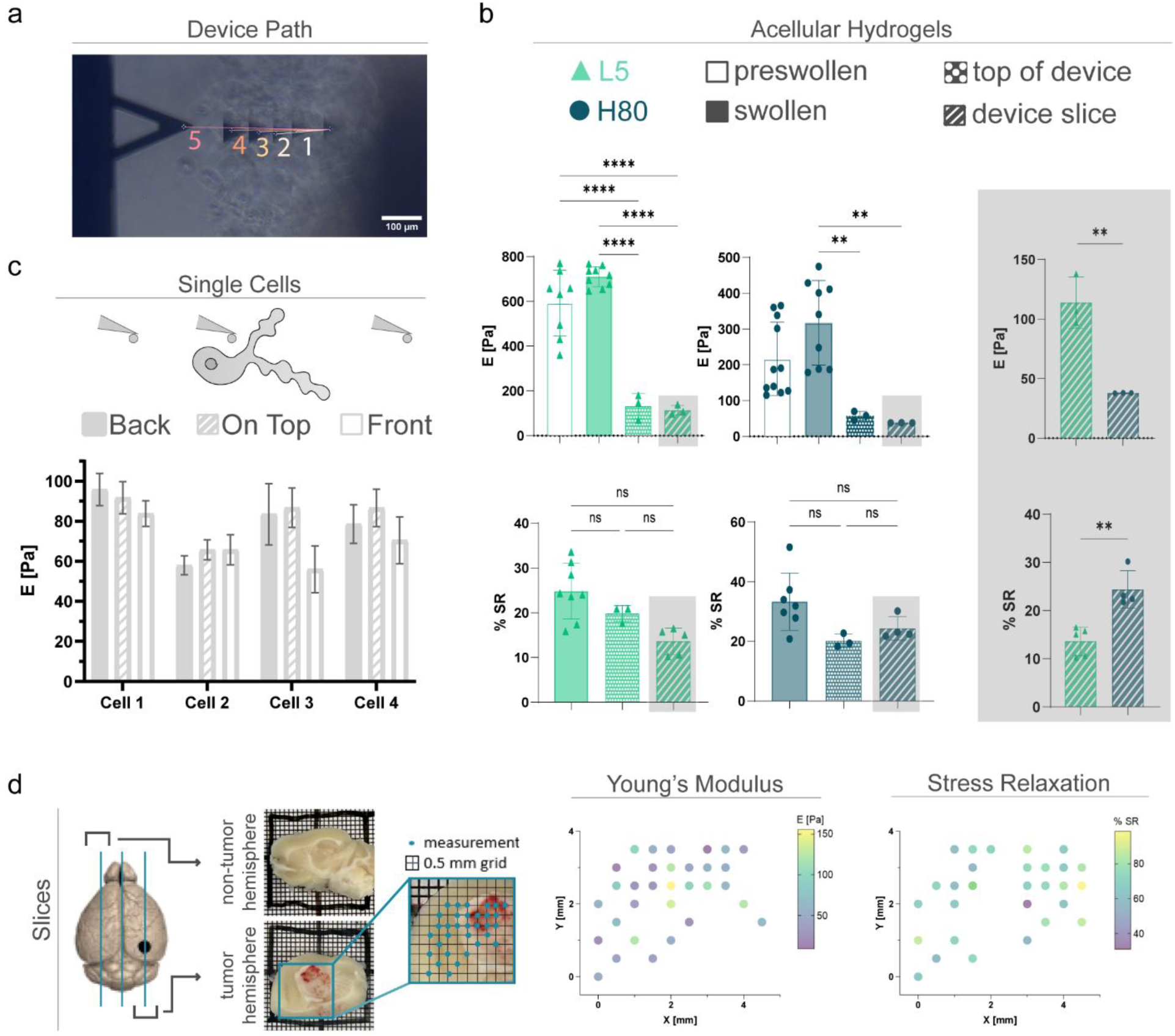
AFM controls. **a.** Quantifying distances between the first and subsequent points in a device path in ImageJ (scale bar 100 μm). **b.** Young’s modulus and stress relaxation measurements for acellular H80 and L5 hydrogels with different swelling and slicing conditions. Each point represents an independent hydrogel. Statistics are obtained by a two-tailed one-way analysis of variance (ANOVA) followed by Tukey’s multiple comparisons test. **c.** Young’s modulus measurements behind, on top, and in front of single cells in H80 hydrogels on day 4. **d.** AFM measurements on tumor and non-tumor mouse brain hemisphere slices. The schematic shows a black point for the tumor cells injection location and blue lines for slicing locations. Representative images of tumor and non-tumor slices on top of a 0.5 mm grid. Inset shows the 0.5 mm grid overlayed on the tumor hemisphere image in Illustrator, with blue dots to signify spatial mapping of measurements. Representative spatial maps of both Young’s modulus and stress relaxation measurements on the tumor hemisphere slice do not show distinct trends.

**Figure S3.**
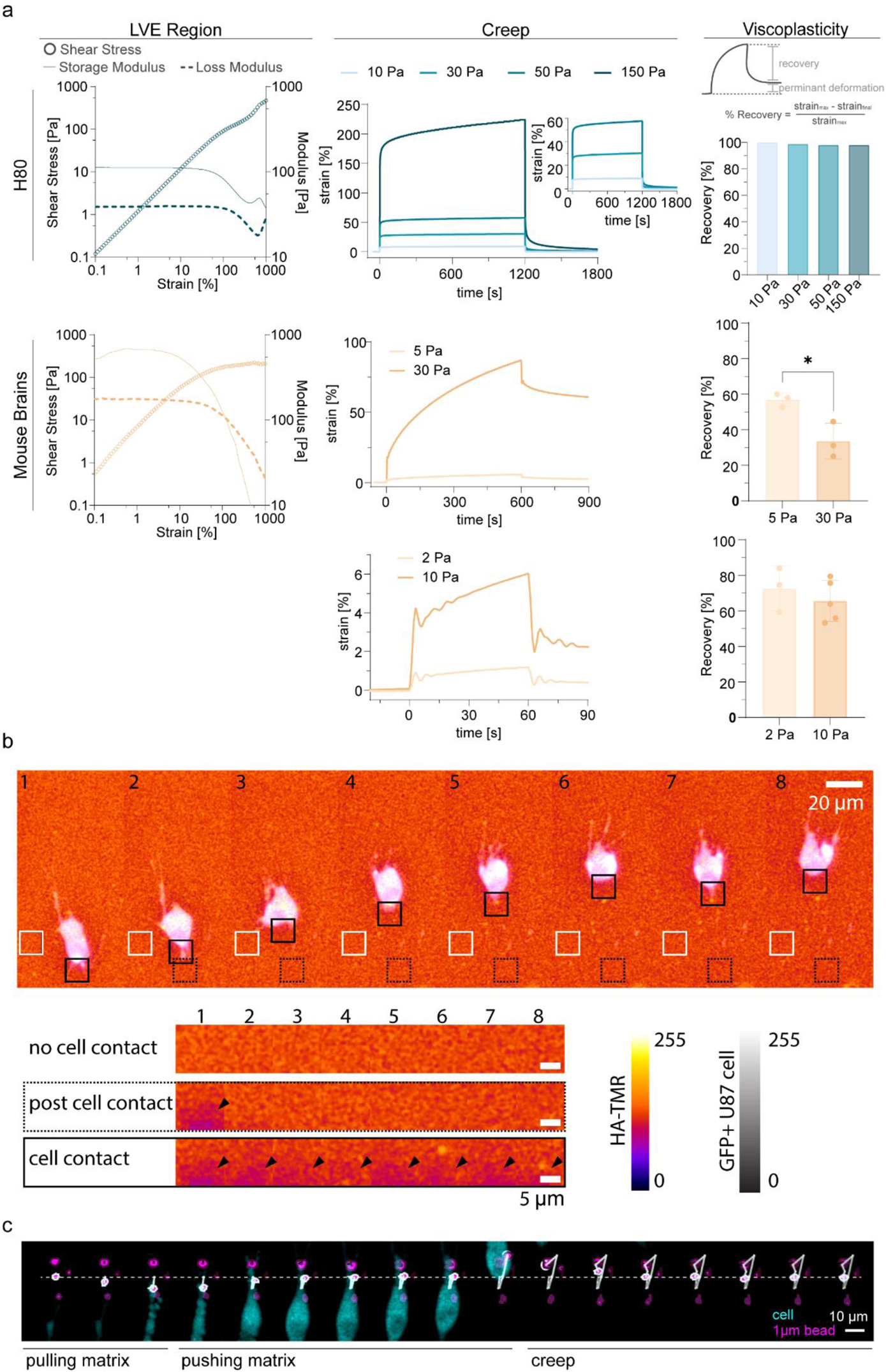
H80 shows little viscoplasticity. **a.** Oscillatory amplitude sweep and creep tests were performed on H80 hydrogels (using a 25 mm 1 angle cone and plate probe) and ∼1mm thick mouse brain samples (using an 8 mm parallel plate probe), where % recovery was plotted for creep tests. Points indicate independent mouse brain samples (n = 3 – 5). **b.** Montage of a GFP+ leader cell invading through a H80 matrix fluorescently tagged with a cysteine-TMR probe (frame = 30 min; scale bar = 20 μm). 3 insets are used to show matrix intensity where there is no cell contact (white solid box), after cell contact (black dotted box), and during cell contact (black solid box) overtime (scale bar = 5 μm). **c.** Montage of a leader cell invading through a 1 μm fluorescent-bead-laden H80 hydrogel at a concentration of 0.05% solid (frame = 30 min). Maroon outline around bead far from cell and white outline around bead that comes in contact with the cell. White solid line shows trajectory of the bead and white dashed line indicates the center line of the bead’s starting position (scale bar = 10 μm).

**Figure S4.**
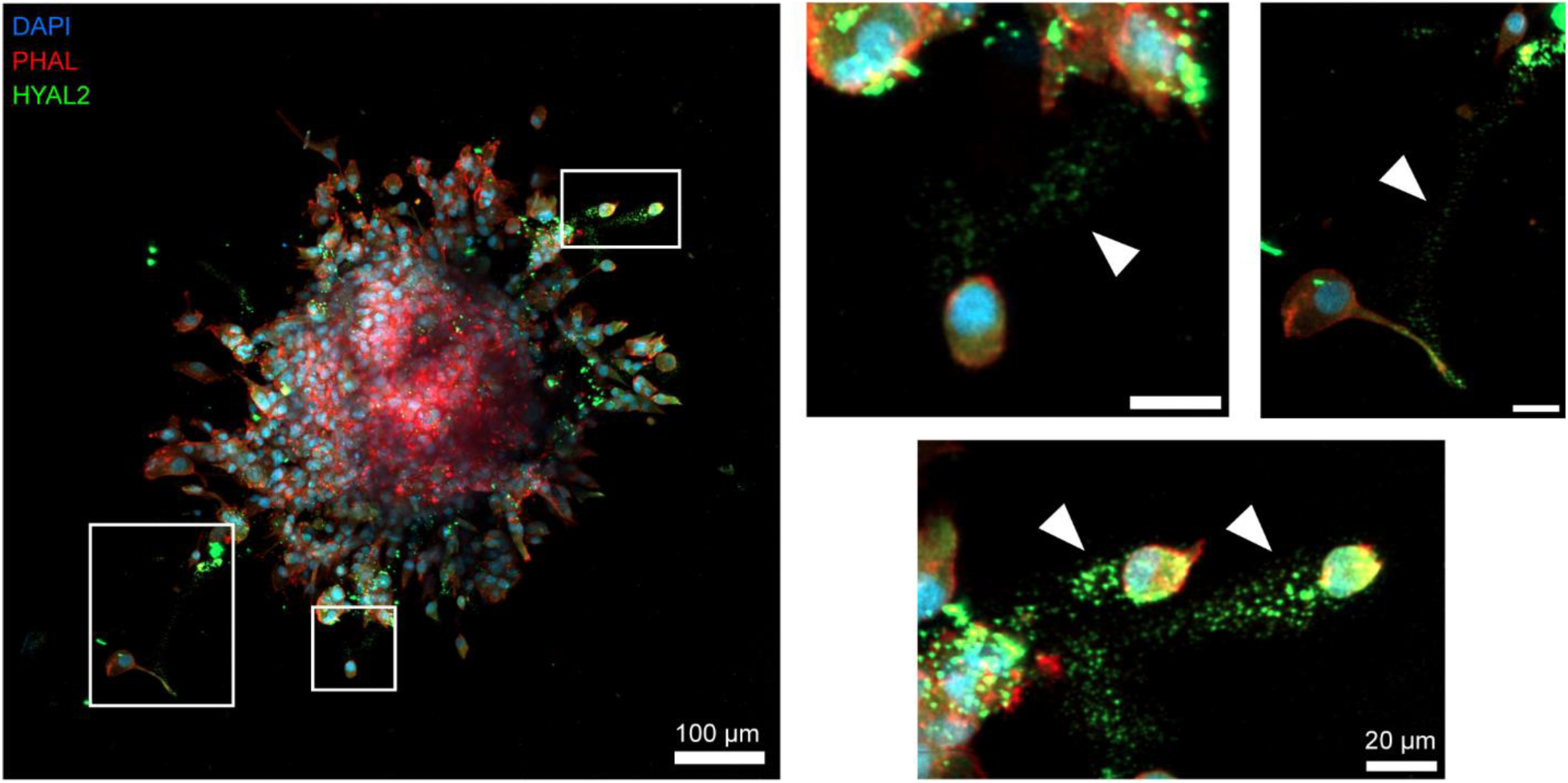
HYAL2 spatial localization in H80 spheroid invasion. Representative confocal z-projections reveal spatial localization of HYAL2 trailing behind leader cells in H80 hydrogels fixed after 2 days (scale bar = 100 μm; insets = 20 μm).

**Figure S5.**
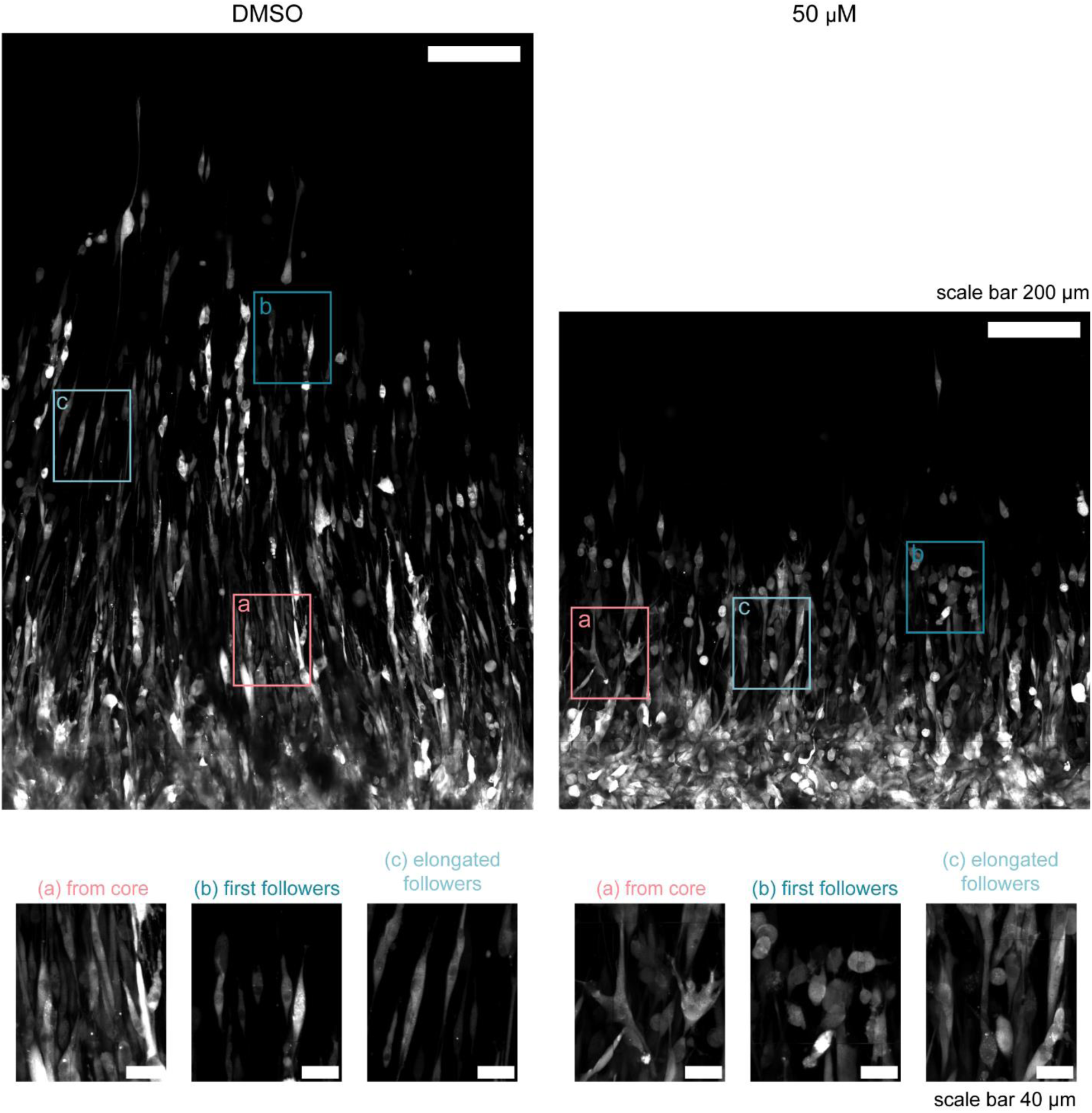
Follower cell alignment. Representative confocal z-projections of the H80 invasive fractions in the DMSO control and 50 μM apigenin conditions taken on day 4 (scale bar = 200 μm; insets = 40 μm). Inset areas display three locations within the invasive fraction with and without substantial cell alignment: *(a) from core* shows cells that are elongated after leaving the core, *(b) first followers* shows elongated cells directly behind the leader cells, and *(c) elongated followers* shows elongated cells within the invasive fraction. While many cells in the DMSO control have an elongated morphology, many cells in the 50 μM take on other morphologies, such as rounded and spread.

**Figure S6.**
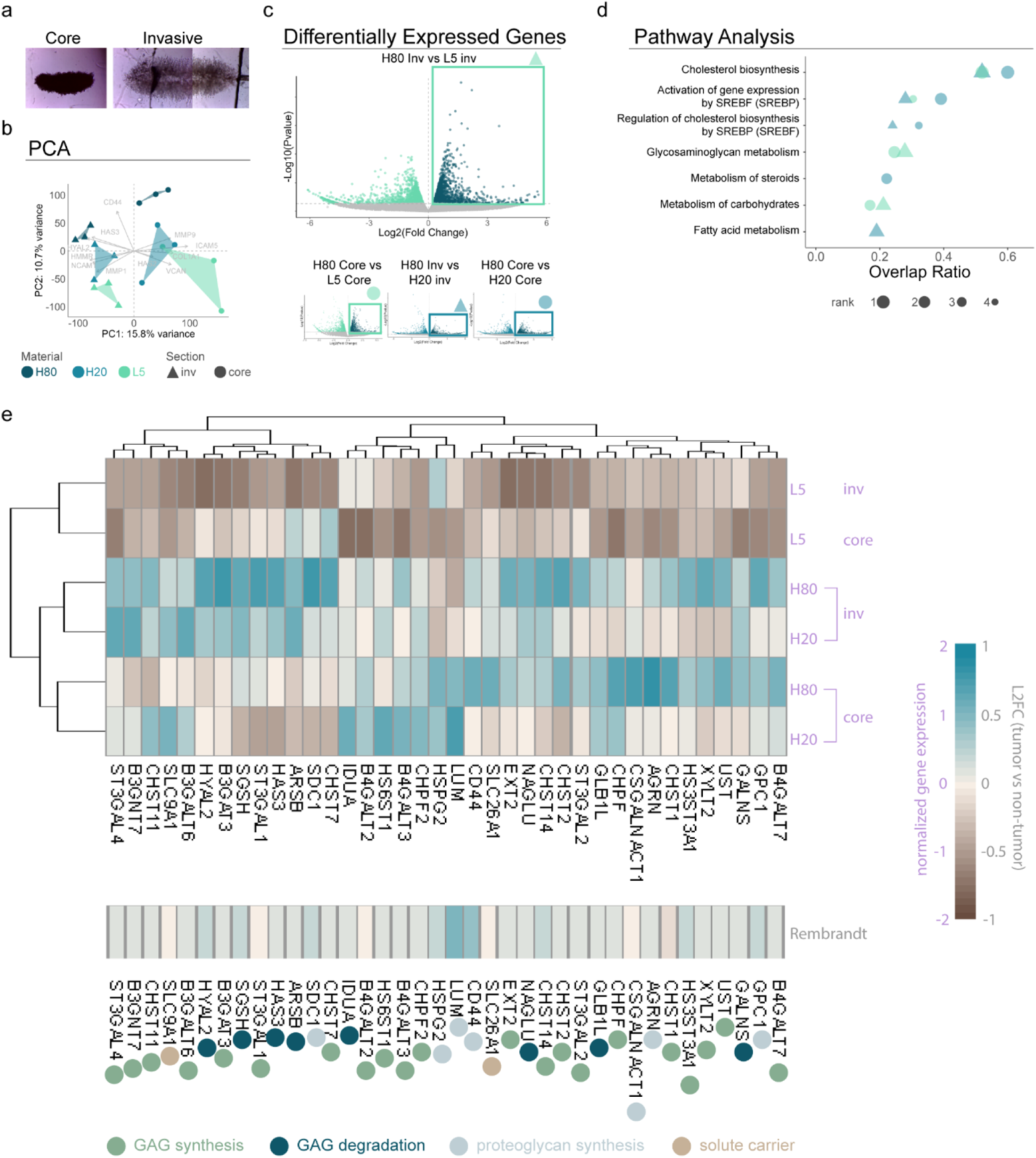
Transcriptomic analysis of H80, H20, and L5 invasive and core samples. **a.** Representative dissection images of core and invasive fractions. **b.** Principal component analysis displays sample clustering within each group. ECM-related eigenvectors are plotted to show genes that are important in core and invasive samples. **c.** Four lists of upregulated differentially expressed genes (DEGs) in H80 compared to its sectioned counterpart in either H20 or L5 are obtained based on a statistical cutoff of |Log2FC| > 0 and p-value < 0.05. **d.** The four DEG lists are run through a pathway analysis using the Reactome (2022) database on Enrichr. Pathways were ranked by their average overlap ratio (DEGs in the pathway: total number of genes in the pathway). Pathways are chosen based on the top 4 pathways for each DEG list based on adjusted p-value (only adjusted p-values < 0.05 are displayed). Dot sizes are chosen based on their adjusted p-values within each list (ranked 1 – 4 with 1 being the smallest adjusted p-value value and largest dot size). **e.** Heat maps of GAG metabolism genes upregulated in H80 fractions, shown as normalized gene expression of 6 sample groups (top, n = 3 per sample group) and the log_2_ fold change (L2FC) of tumor vs non-tumor samples in the Rembrandt data set (bottom). Genes are categorized by function in GAG synthesis, GAG degradation, proteoglycan synthesis, or as solute carriers.

**Figure S7.**
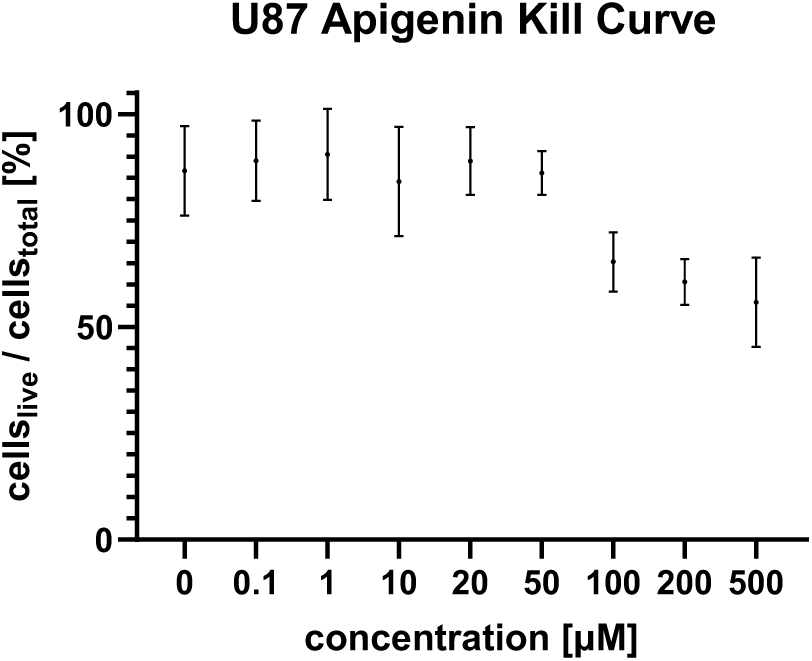
Apigenin cell viability assay. Single-cell laden hydrogels were exposed to varying levels of apigenin, and a live/dead assay was performed after 2 days in culture to assess max concentration of apigenin for cell culture experiments.

